# White-tailed deer (*Odocoileus virginianus*) may serve as a wildlife reservoir for nearly extinct SARS-CoV-2 variants of concern

**DOI:** 10.1101/2022.09.02.506368

**Authors:** Leonardo C. Caserta, Mathias Martins, Salman L. Butt, Nicholas A. Hollingshead, Lina M. Covaleda, Sohel Ahmed, Mia Everts, Krysten L. Schuler, Diego G. Diel

## Abstract

The spillover of severe acute respiratory syndrome coronavirus 2 (SARS-CoV-2) from humans into white-tailed deer (WTD) and its ability to transmit from deer-to-deer raised concerns about the role of WTD in the epidemiology and ecology of the virus. In the present study, we conducted a comprehensive investigation to assess the prevalence, genetic diversity, and evolution of SARS-CoV-2 in WTD in the State of New York (NY). A total of 5,462 retropharyngeal lymph node (RPLN) samples collected from free-ranging hunter-harvested WTD during the hunting seasons of 2020 (Season 1, September-December 2020, n=2,700) and 2021 (Season 2, September-December 2021, n=2,762) were tested by SARS-CoV-2 real-time RT-PCR. SARS-CoV-2 RNA was detected in 17 samples (0.6%) from Season 1 and in 583 (21.1%) samples from Season 2. Hotspots of infection were identified in multiple confined geographic areas of NY. Sequence analysis of SARS-CoV-2 genomes from 164 samples demonstrated the presence multipls SARS-CoV-2 lineages as well as the co-circulation of three major variants of concern (VOCs) (Alpha, Gamma, and Delta) in WTD. Our analysis suggests the occurrence of multiple spillover events (human-to-deer) of the Alpha and Delta lineages with subsequent deer-to-deer transmission of the viruses. Detection of Alpha and Gamma variants in WTD long after their broad circulation in humans in NY suggests that WTD may serve as a wildlife reservoir for VOCs no longer circulating in humans. Thus, implementation of continuous surveillance programs to monitor SARS-CoV-2 dynamics in WTD are warranted, and measures to minimize virus transmission between humans and animals are urgently needed.

**SIGNIFICANCE:** White-tailed deer (WTD) are highly susceptible to severe acute respiratory syndrome coronavirus 2 (SARS-CoV-2) and are known to efficiently transmit the virus to other susceptible animals. Evidence of natural exposure or infection of wild WTD in North America raised significant concerns about their role on the ecology of the virus and its impact on the control of the coronavirus disease 2019 (COVID-19) pandemic. This comprehensive study demonstrates widespread infection of SARS-CoV-2 in the WTD populations across the State of New York. Additionally, we showed co-circulation of three major SARS-CoV-2 variants of concern (VOCs) in this wildlife population, long after their broad circulation in humans. These findings indicate that WTD – the most abundant large mammal in North America – may serve as a reservoir for variant SARS-CoV-2 strains that no longer circulate in the human population.

## INTRODUCTION

The coronavirus disease 2019 (COVID-19) was declared a pandemic in March of 2020, and as of August 2022 has incurred over 590 million human cases and more than 6.4 million deaths globally (1). COVID-19 is caused by severe acute respiratory syndrome coronavirus 2 (SARS-CoV-2), a new zoonotic virus for which most of the first known human infections were linked to the Huanan Seafood Wholesale market in Wuhan, China, where several live wild animal species were sold (2). SARS-CoV-2 is a single-stranded RNA virus within the *Sarbecovirus* subgenus, *Betacoronavirus* genus, of the family *Coronaviridae* (3). Analysis of the genome sequence of SARS-CoV-2 revealed high similarity to coronaviruses circulating in bats in China, suggesting that bats are the most likely source of the ancestral virus that originated SARS-CoV-2 (4). While the closest bat coronaviruses (RaTG13, RmYN02, RpYN06 and PrC31) are phylogenetically related to SARS-CoV-2, they present several mutations across the genome that distinguish them from SARS-CoV-2 indicating that direct transmission of the virus from bats to humans was unlikely (4, 5). These observations point to the involvement of a yet unidentified animal species that served as an intermediate host and enabled spillover of the virus into humans (6).

Comparative sequence analyses of the main cellular receptor for SARS-CoV-2, the angiotensin converting enzyme 2 (ACE2), from more than 400 animal species suggested a broad host range for the virus (7). Notably, the ACE2 of three species of deer, including reindeer (*Rangifer tarandus*), Père David’s deer (*Elaphurus davidianus*) and white-tailed deer - WTD (*Odocoileus virginianus*) share a high similarity to the human ACE2 and were predicted to allow binding and entry of SARS-CoV-2 into deer cells (7). We and others have confirmed these *in silico* predictions and demonstrated that WTD are highly susceptible to SARS-CoV-2 infection (8, 9). Most importantly, intranasal inoculation of SARS-CoV-2 in WTD resulted in virus replication and shedding, which led to efficient deer-to-deer transmission of the virus (8–10). These findings placed WTD - a species broadly distributed in North America with an estimated population size of 30 million animals (11) – at the center of investigations which demonstrated exposure of free ranging WTD to SARS-CoV-2 and highlighted their potential to serve as a reservoir for SARS-CoV-2 in North America. Notably, results from the first studies revealed prevalence rates varying from ∼30-40%, with animals testing positive for SARS-CoV-2 antibodies and/or viral RNA (12–17). Importantly, detection of SARS-CoV-2 RNA in respiratory secretions and tissues from several of the sampled animals, suggested recent infection with the virus (12–15, 17). Additionally, analysis of SARS-CoV-2 sequences recovered from WTD revealed multiple introductions from humans and suggested onward transmission of the virus in free-ranging deer populations (12–15). The pathways of human-to-deer transmission of SARS-CoV-2, however, are not yet understood. These findings highlight the need to establish surveillance programs that will allow continuous monitoring of the circulation, distribution, and evolution of SARS-CoV-2 in WTD populations.

In this study, we conducted a comprehensive investigation to assess the prevalence, genetic diversity, and evolution of SARS-CoV-2 in WTD in New York (NY) State. All samples included in our study (n = 5462) consisted of retropharyngeal lymph nodes (RPLN) – one of the major sites of SARS-CoV-2 replication in WTD (10) – that were collected as part of the New York State’s Chronic Wasting Disease (CWD) Surveillance program during two hunting/CWD testing seasons (Season 1, 2020; and Season 2, 2021).

## RESULTS

### Prevalence of SARS-CoV-2 infection in white-tailed deer in New York

A total of 5,462 RPLN samples collected as part of the New York State’s CWD Surveillance program from wild hunter-harvested WTD during the hunting seasons of 2020 (Season 1, September 2020 through December 2020) and 2021 (Season 2, July of 2021 through December 2021) were tested for SARS-CoV-2 by real-time RT-PCR. In Season 1, positive samples were detected in October and November 2020, whereas in Season 2 positive cases were detected between October and December 2021, with peak positivity being detected between November 1 and 15, 2021 (Fig. 1A). SARS-CoV-2 RNA was detected in 17 of 2,700 (0.6%) samples from Season 1 and in 583 of 2,762 (21.1%) samples collected on Season 2 (Fig. 1B). Viral RNA loads, based on RT-PCR cycle threshold values (Cts), varied slightly between the seasons (mean Ct values for Season 1 was 29.8 ± 3.4 [mean ± SD, range from 24 to 37], whereas for Season 2 mean Ct was 30.8 ± 3.4 [range from 21 to 37]) (Fig. 1C), however, no statistical differences were observed in Ct values between the two Seasons (*p >* 0.2). All RPLN samples with Ct values lower than 30 (11 samples from Season 1, and 205 samples from Season 2), were subjected to virus isolation in Vero-E6 TMPRRSS2 cells (10, 18). Infectious virus was recovered from seven samples collected in Season 2 (Fig. 1D). Together, these results showed a significant increase (35-fold, chi-square = 585.5, P < 0.0001) in the prevalence of SARS-CoV-2 infection between the 2020 and 2021 seasons and demonstrate a broad circulation of the virus in WTD in NY.

**FIG 1.**
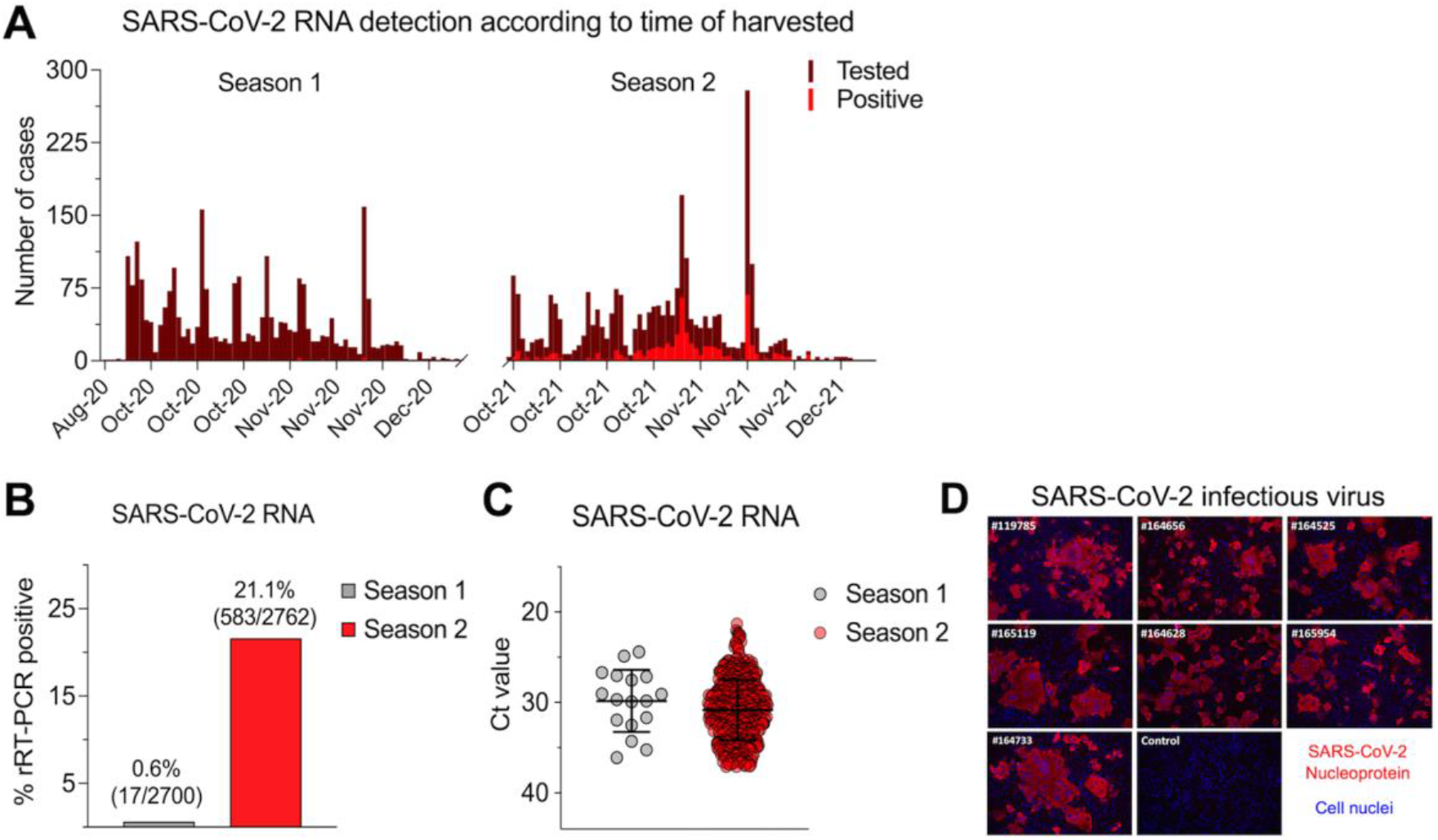
Prevalence of SARS-CoV-2 infection in WTD in NY. A total of 5,462 retropharyngeal lymph node (RPLN) samples were collected from free-ranging hunter-harvested white-tailed deer (WTD) during the hunting Seasons 1 (September - December 2020, n = 2,700) and Season 2 (September - December 2021, n = 2,762) in the State of New York (NY). (A) RPLN tested for the presence of SARS-CoV-2 viral RNA using real-time reverse transcriptase PCR (rRT–PCR). The number of samples tested and SARS-COV-2 positive samples in Season 1 and Season 2 is shown in the graphic. (B) Distribution of positive samples per season. Number of SARS-COV-2 RNA positive RPLN from WTD. (C) SARS-CoV-2 RNA detection by rRT–PCR presented as cycle threshold (*Ct*s) value. All the Ct values for positive samples from season 1 (*n* = 17) and season 2 (*n* = 583) are shown. (D) Infectious SARS-CoV-2 recovered from seven samples (sample #IDs are indicated in white). Virus isolation was confirmed by immunofluorescence assay using a monoclonal antibody anti-SARS-CoV-2-nucleoprotein (N) (red). Nuclear counterstain was performed with DAPI (blue). 20x magnification.

### Hotspots of SARS-CoV-2 infection among white-tailed deer in the State of New York

The samples tested in our study were collected statewide from WTD harvested in 57 of 62 counties in New York. The geographic distribution of the samples from Season 1 and Season 2 and those that tested positive for SARS-CoV-2 is presented in Fig. 2A. In Season 1, 17 positive samples were detected in ten of the 62 counties in the state including Orleans (n = 1) and Monroe County (n = 2) in the Finger Lakes region, Steuben (n = 4) and Chemung County (n = 1) in the Southern Tier, Delaware (n = 1), Sullivan (n = 3) and Ulster (n = 2) in the Hudson Valley Region, Columbia (n = 1) and Greene County (n = 1) in the Capital Region, and Clinton County (n = 1) in the North Country (Fig. 2A). In Season 2, the increased prevalence of SARS-CoV-2 infection in WTD was accompanied by a marked increase in the geographic distribution of the virus with positive cases being detected in 48 counties (Fig. 2A and B). The proportion of positive samples detected in each county is presented in Fig. 2B. Positive SARS-CoV-2 cases were detected in nine of ten geographic regions of the state (Western New York, Finger Lakes, Southern Tier, Central New York, North Country, Mohawk Valley, Capital Region, Mid-Hudson, and Long Island) (Fig. 2A). No samples were collected in New York City. The counties with the highest number of cases were Allegany (n = 72), Cattaraugus (n = 32), and Steuben (n = 31) in the Southern Tier region, Chautauqua (n = 62) in Western New York region, and Orange (n = 32) and Sullivan (n = 44) in the Hudson Valley region. Together these counties accounted for 47% of the cases detected across the state (n = 273). A summary of the samples tested, and SARS-CoV-2 positivity detected by county is presented in Table S1.

**FIG 2.**
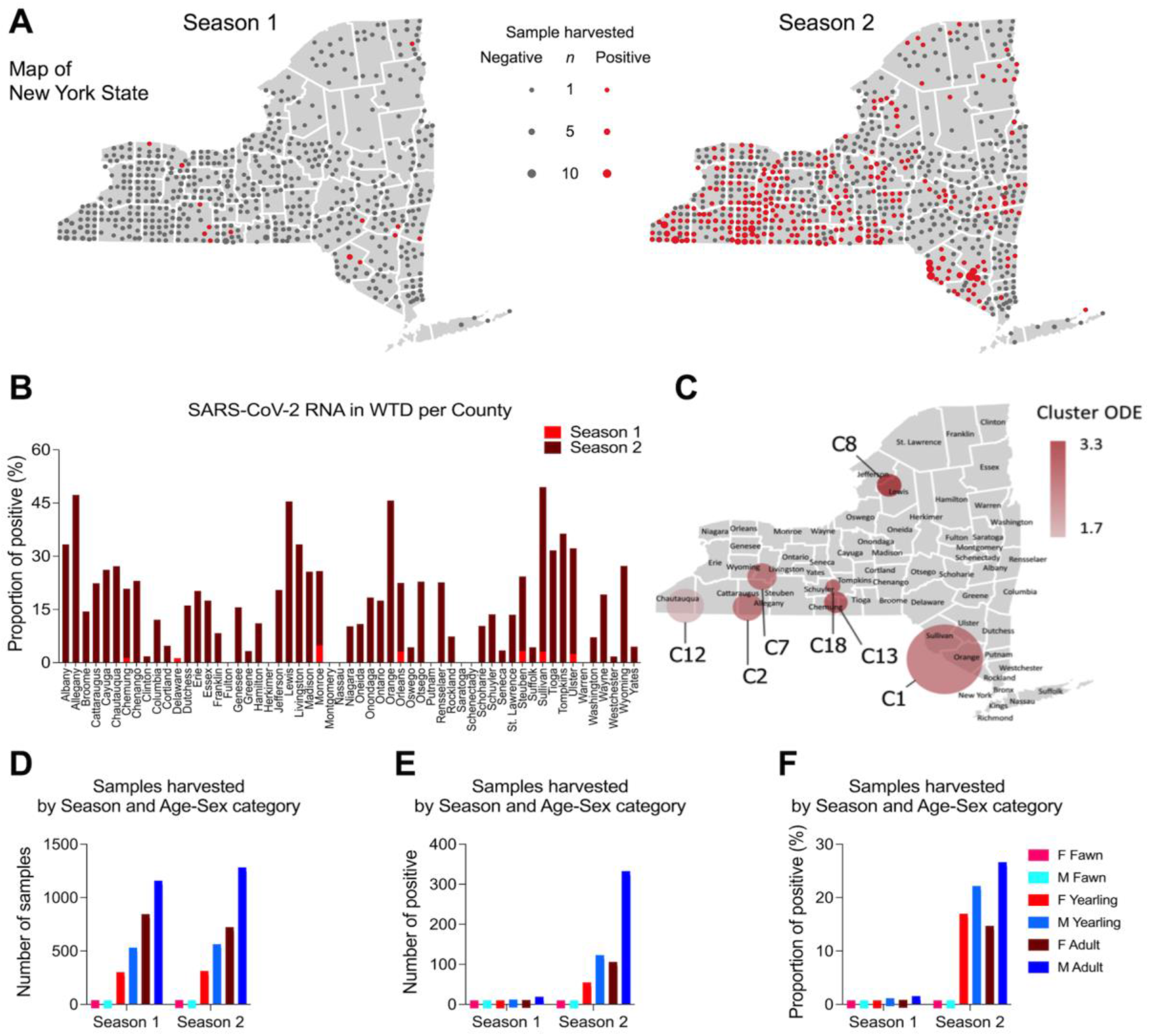
Demographics of WTD population sampled and tested for SARS-CoV-2. Retropharyngeal lymph node (RPLN) samples were collected from free-ranging hunter-harvested white-tailed deer (WTD) during the hunting Seasons 1 (September - December 2020, n = 2,700) and Season 2 (September - December 2021, n = 2,762) in the State of New York (NY). (A) Sampling distribution and positivity across the State of New York. SARS-CoV-2 RNA was detected in 17 samples (0.6%) from Season 1 and in 583 (21.1%) samples from Season 2. (B) Proportion of SARS-CoV-2 positive samples in Season 1 and Season 2. (C) High-risk areas or hotspots of SARS-CoV-2 infection in WTD in New York. Spatial cluster analysis performed with the samples from Season 2 (*n* = 2,762). Nineteen spatial clusters (C1 - C19) containing from 3 to 57 positive samples collected within a radius of 10.6 to 55 kilometers (Km) from each other were identified. Seven hotspots with high-risk for SARS-CoV-2 infection in WTD (relative risk [RR] > 1.76) are highlighted in the map. (D) Total number of collected samples based on sex and age of WTD during Season 1 and Season 2. Animals were distributed in three age groups as follows: < 1.5 years old (fawns), ≥ 1.5 and < 2.5 years old (yearlings), ≥ 2.5 years old (adults). (E) Number of SARS-CoV-2 positive samples based on sex and age of WTD during Season 1 and Season 2. (F) Proportion of positive samples based on sex and age of WTD during Season 1 and Season 2.

To identify high-risk areas or hotspots of SARS-CoV-2 infection in WTD in New York, spatial cluster analysis was performed with the samples from Season 2. This analysis included all 2,762 samples tested in Season 2 and relied on a likelihood ratio test and explored the maximum number of positive SARS-CoV-2 cases that were detected in a given geographic location. Nineteen spatial clusters (C1 - C19) containing 3 to 57 positive samples were identified (Table S2). These SARS-CoV-2 positive WTD clusters comprised samples collected within a radius of 10.6 to 80 kilometers (Km) from each other (Table S2). Among the 19 clusters identified, seven hotspots with high-risk for SARS-CoV-2 infection in WTD (relative risk [RR] > 1.76) were detected (Fig. 2C; Table S2), most of them (C2, C7, C13 and C18) in the Southern Tier region of NY. One cluster (C12, RR = 1.76) was formed by samples collected in Chautauqua Western New York Region and a small cluster (C8, RR = 3.38) in the North Country regions of the state, respectively (Fig. 2C). From the Southern Tier clusters, Cluster 2 (RR = 2.7) involved two counties (Allegany and Cattaraugus), Clusters 13 (RR = 2.85), Cluster 7 (RR = 2.63) and Cluster 18 (RR = 2.74) comprised one county each. The larger cluster identified in our analysis, Cluster 1 at Hudson Valley (n = 89 cases) involved three counties (Sullivan, Orange, and Ulster). These results revealed several hotspots of SARS-CoV-2 infection in WTD in NY, most of which overlap with geographic areas with the highest deer population densities and deer harvest rates in the state.

### Male white-tailed deer are at a higher risk of infection with SARS-CoV-2

The distribution of positive SARS-CoV-2 cases based on sex and age of the animals sampled in our study was also investigated. Sex and age of all WTD harvested from Season 1 and Season 2 were determined and all animals were distributed in three age groups as follows: < 1.5 years old (fawns), ≥ 1.5 and < 2.5 years old (yearlings), and ≥ 2.5 years old (adults). Overall, the number of males in each age group sampled during Season 1 and Season 2 were slightly higher than the number of females (Fig. 2D), with the number of males that tested positive for SARS-CoV-2 in all age groups being 2-3 times higher than the number of females in the corresponding age group (Fig. 2E). As shown in Fig. 2F the proportion of males that tested positive in all age groups was also markedly higher than the proportion of positive females detected. Logistic regression analysis confirmed that males were more likely to test positive for SARS-CoV-2 than females (OR = 1.952, 95% CI = 1.591 - 2.407), with adult males being more likely to become infected with SARS-CoV-2 than younger yearling males (OR = 1.906, 95% CI = 1.55 - 2.35) (Table S3). These results suggest that adult male WTD are at a higher risk of infection with SARS-CoV-2.

### Co-circulation of multiple SARS-CoV-2 variants of concern in white-tailed deer

To assess the genetic makeup and diversity of SARS-CoV-2 in WTD, we performed whole genome sequencing on 216 samples (including 11 samples from Season 1 and 205 samples from Season 2) that tested positive for SARS-CoV-2 and presented a RT-PCR Ct value < 30. Consensus genome sequences of 216 samples were assembled and subjected to SARS-CoV-2 lineage classification using Pangolin version 4.0.6 (19). Of the 216 samples sequenced, 164 genomes passed the Pangolin QC thresholds for minimum length and maximum N content (10%) and were used for downstream analyses. A total of nine SARS-CoV-2 genomes were recovered from samples from Season 1, and 155 from samples from Season 2. The geographic distribution of the samples from which complete or near complete SARS-CoV-2 genome sequences were obtained according to the season is presented in Fig. 3A. Nine lineages including three major variants of concern (VOCs) (Alpha [B.1.1.7], Gamma [B.1.1.28, P.1] and Delta [B.1.617.2]) were detected in WTD samples in NY. The SARS-CoV-2 sequences recovered in Season 1 were classified as B.1, B.1.1, B.1.2, B1.243, B.1.409, B.1.507 and Alpha, whereas in Season 2, B.1.1, B.1.2, B.1.517, Alpha, Gamma and Delta lineages were detected. The geographic distribution of these lineages across the state is presented in Fig. 3B. Most of the samples sequenced in our study were classified within one of the three majors VOCs identified Alpha, Gamma, or Delta. Interestingly, Alpha and Delta variants were detected in several geographic regions throughout the state, while the samples classified as Gamma (n = 27) formed a localized cluster in the Southern Tier County of Allegany (Fig. 3B). Notably, there was one sample detected in Season 1 (161392; GISAID EPI_ISL_13610582 and GenBank OP006342) that was classified by Pangolin as B.1.617.2 (Delta), but the lineage for this sample was unassigned by Scorpio (19). Analysis of the mutation profile of the sample revealed the presence of only 8 of 20 B.1.617.2 lineage defining mutations (ORF1b, P314L; S, T19R, T478K, and D614G; ORF7a, T120I; and N D63G, R203M and D377Y), thus a lineage was not assigned to this sample.

**FIG 3.**
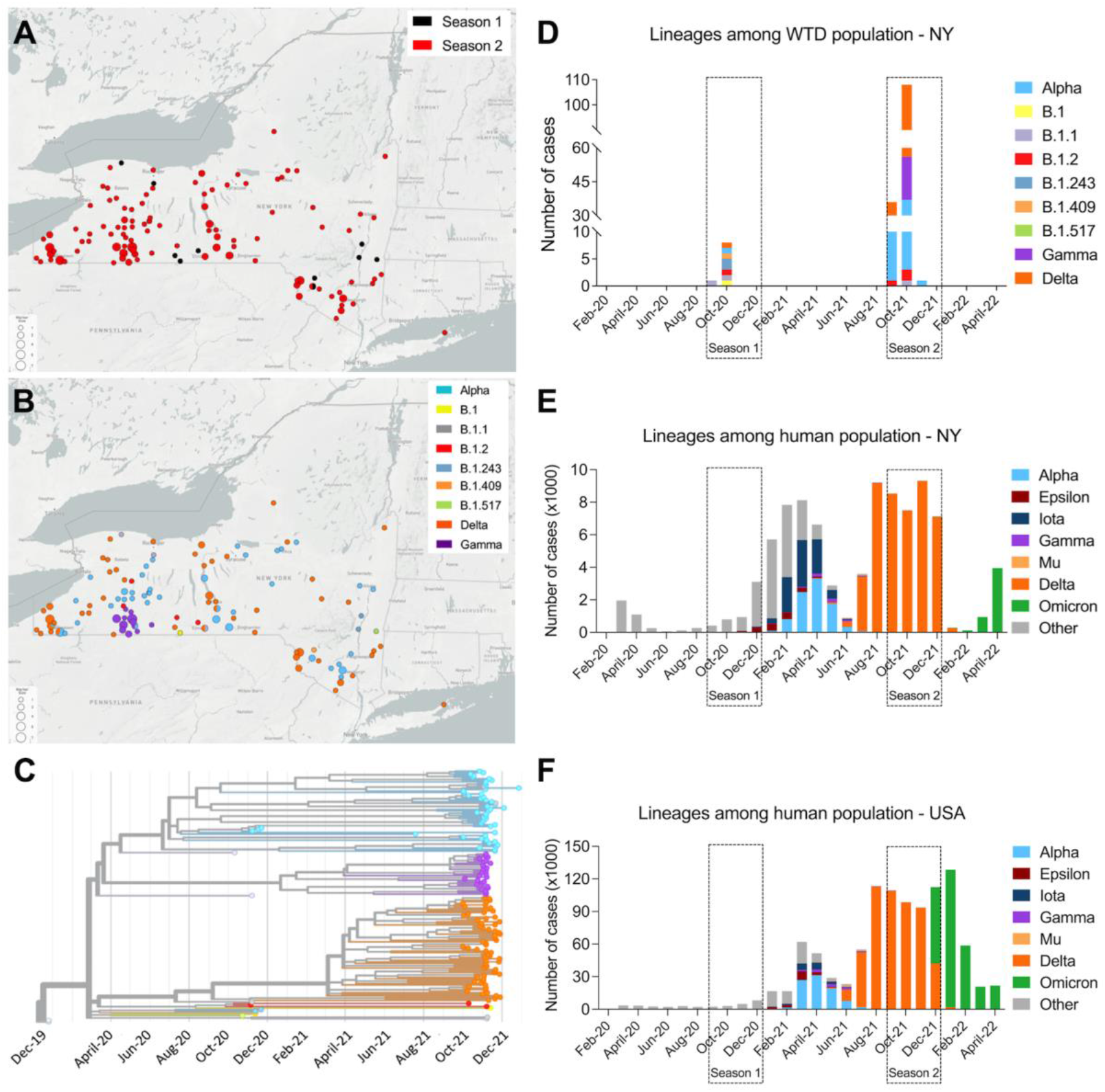
Distribution of SARS-CoV-2 lineages across the State of New York (NY) and phylogenetic relatedness of viral genomes. (A) Geographic distribution of the 164 samples from which complete or near complete SARS-CoV-2 genome sequences were obtained and according to the season is presented (9 and 155 genomes from Season 1 and Season 2, respectively). (B) Geographic distribution of the SARS-CoV-2 lineages detected in free-ranging white-tailed deer (WTD) in this study. (C) Distribution of SARS-CoV-2 lineages among WTD population in NY during Season 1 and Season 2. (D) Phylogenetic tree showing 164 SARS-CoV-2 genomes obtained from WTD in New York across time. (E) Monthly distribution of SARS-CoV-2 human cases (February 2020 to May 2022) from NY and (F) United States of America (US). The colored stacked bars represent proportion of each SARS-CoV-2 lineages.

Phylogenetic analysis performed with the SARS-CoV-2 sequences revealed three large clusters corresponding to three SARS-CoV-2 VOCs (Alpha, Gamma, and Delta) identified in the sampled WTD population (Fig. 3C). Together these results demonstrate the concomitant circulation of Alpha, Gamma, and Delta VOCs in WTD in NY in 2021.

Given that most of the samples that were identified as Alpha, Gamma or Delta variants in WTD in our study were detected between October and December 2021 (Season 2), we analyzed the circulation of these VOCs in humans. The number of human SARS-CoV-2 cases per lineage based on available sequencing information for the State of NY and the US over a time frame (February 2020 to April 2022) overlapping the WTD sampling performed in this study. This data clearly shows an overlap in detection of the Delta variant in humans and in WTD in NY and across the US (Fig. 3D, E and F). Notably, while the Alpha and Gamma variants were circulating in WTD in NY in November and December 2021 (Fig. 3D), detection of these VOCs in humans peaked between April and June, 2021, with only sporadic detections after August (Fig. 3E). Three Alpha sequences were reported in NY in September 2021 based on data available at GISAID, with a single last detection occurring in February 2022, while the two last Gamma sequences were detected in October 2021. All these latest Alpha and Gamma detections in humans in the state occurred in New York City, which was not included in the WTD sampling in this study. Interestingly, the Alpha variant was detected in multiple geographic locations in WTD across NY, in a time where there is no evidence that the virus was broadly circulating in humans in the state. The time lapse between the peak of detection of Alpha and Gamma variants in humans and in WTD, suggest that these variants may have become established in WTD in NY.

### Phylogenomic analysis of SARS-CoV-2 variants detected in white-tail deer

The phylogenetic relationship of the WTD SARS-CoV-2 samples characterized in our study (n = 164) were compared to other sequences derived from WTD (n = 159) available in the EpiCoV database in Global Initiative on Sharing All Influenza Data (GISAID). As of August 2022, most available SARS-CoV-2 sequences in GISAID were classified as B.1-like or Delta lineages, whereas most sequences obtained in this study were classified as Delta variant lineage, followed by Alpha and Gamma lineages, respectively. Within each of the major lineages identified in this study (Alpha, Gamma and Delta), samples obtained from close geographic locations (e.g. same county, or neighboring counties) formed clusters of closely related SARS-CoV-2 sequences (Fig. 4). Interestingly, the closest phylogenetic relationship was observed among the Gamma sequences, with all sequences obtained forming a monophyletic cluster (Fig. 4). Pairwise nucleotide analysis demonstrated that the Gamma lineage sequences share up to 99.99 % similarity. All Gamma sequences were obtained from WTD harvested in 6-25 km from each other, within Allegany County. Consistent with the broader geographic distribution throughout state (Fig. 3C), the Alpha and Delta lineage sequences formed multiple smaller clusters, with the Delta sequences from NY being interspersed with Delta variant clusters detected in WTD in other states of the US (Fig. 4).

**FIG 4.**
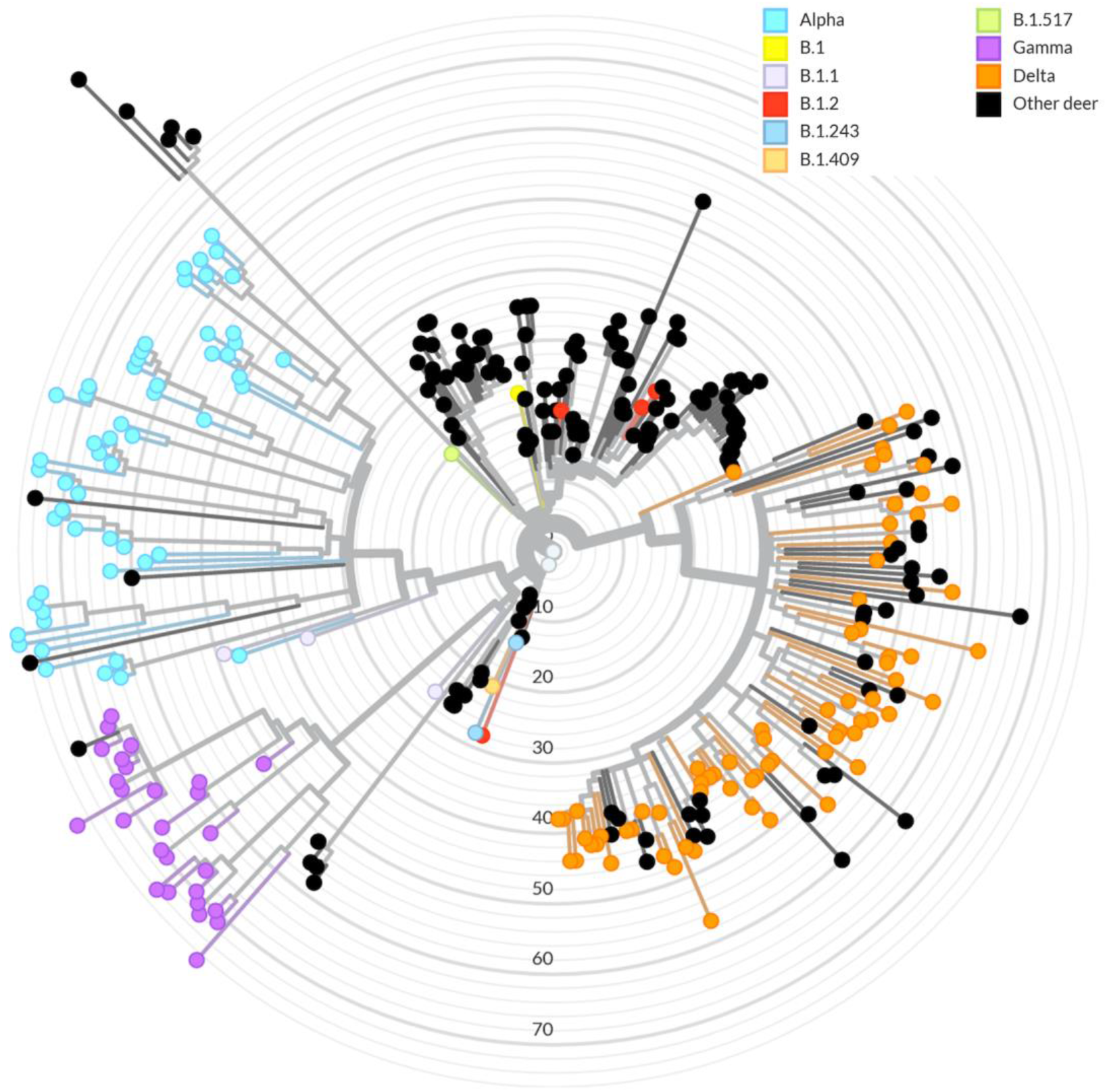
Phylogenomic analysis of SARS-CoV-2 derived from WTD. Phylogeny of 164 SARS-CoV-2 sequences derived from white-tail deer (WTD) in the State of New York during Season 1 and Season 2 in context of 159 SARS-CoV-2 sequences derived from WTD available in GISAID. The node colors (sky blue: Alpha; purple: Gamma and orange: Delta) represent SARS-CoV2 variants of concern (VOCs) derived from WTD sequences from this study and the black nodes represent the Pan-US SARS-CoV-2 sequences derived from WTD.

### Evolution and mutation profile of SARS-CoV-2 in white-tailed deer

The genetic relationship of the WTD SARS-CoV-2 sequences with human SARS-CoV-2 sequences available in GISAID was also investigated. A phylogenetic analysis including all 164 sequences characterized in this study, plus 159 WTD SARS-COV-2 sequences and 3,837 human SARS-CoV-2 genomes available in GISAID was performed (Fig. 5A). The human sequences included in our analysis were obtained between March 5, 2020 and January 30, 2022. No direct links between SARS-CoV-2 sequences detected in WTD and in humans in NYwere observed. Interestingly, WTD sequences from the Alpha (Fig. 5B) and Delta lineages (Fig. 5D) from NY formed 11 and 27 divergent phylogenetic branches, respectively that were interspersed among the human sequences included in our analysis. All Gamma WTD sequences detected here formed a monophyletic group highly divergent from the human derived Gamma lineage sequences (Fig. 5A and C).

**FIG 5.**
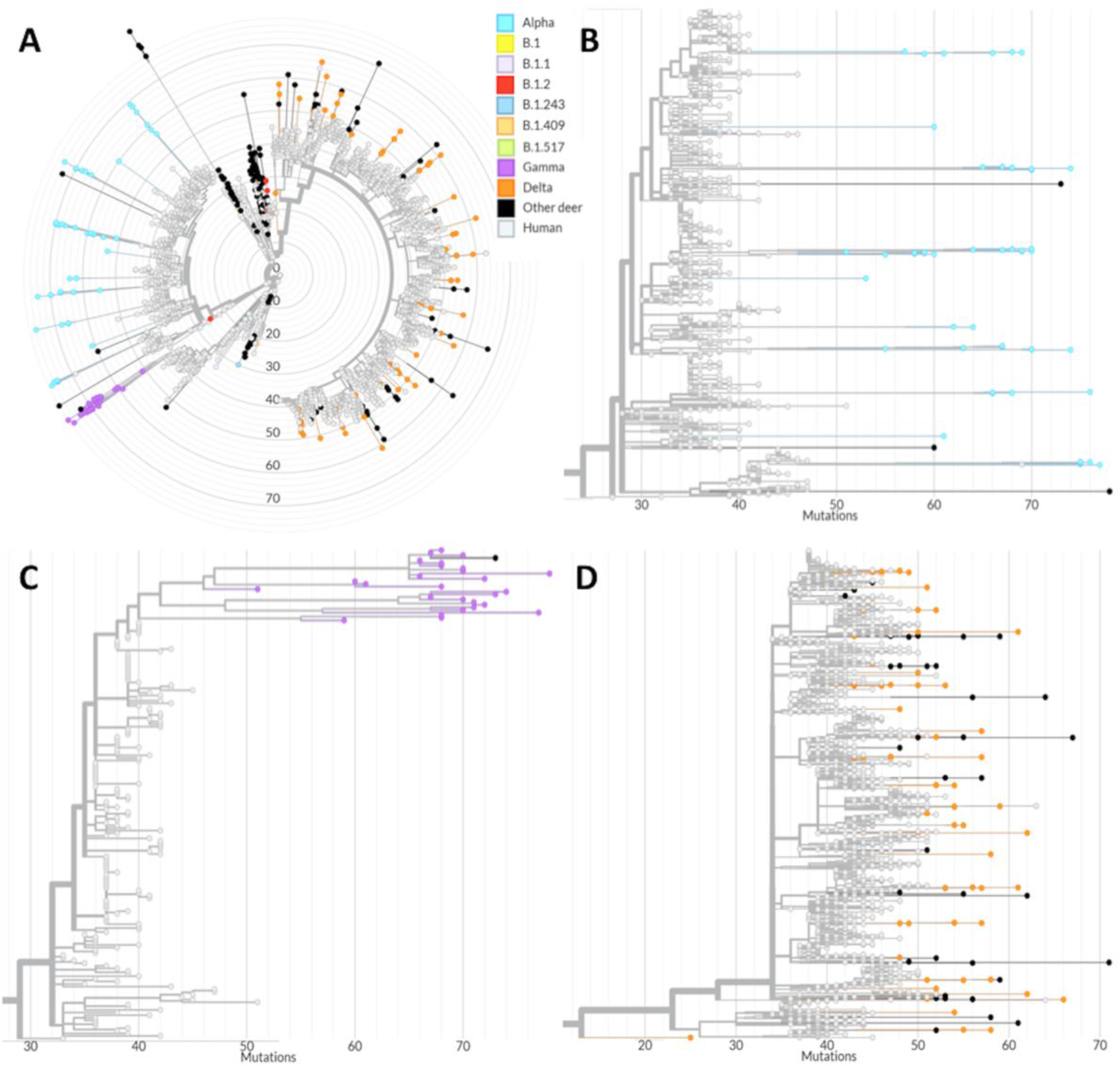
Phylogenomic analysis of SARS-CoV-2 derived from humans and WTD. Divergence between white-tailed deer (WTD) and human derived SARS-CoV-2 sequences. The node colors (sky blue: Alpha; purple: Gamma and orange: Delta) represent SARS-CoV2 variants of concern (VOCs) derived from WTD from this study, black nodes represent the Pan-US SARS-CoV-2 sequences derived from WTD and grey nodes represent SARS-CoV-2 sequences derived from human. The X axis shows the number of mutations compared to the reference sequence Wuhan-1 (GenBank accession number MN908947.3). (A) Phylogenetic tree comprising 164 WTD derived sequences and 3837 human derived sequences. Magnification of the Alpha (B), Gamma (C), and Delta (D) clusters.

Phylogenetic analysis of SARS-CoV-2 detected in WTD revealed a high genetic diversity and marked evolution of the viruses detected in this animal species when compared to human derived SARS-CoV-2 sequences. The WTD SARS-CoV-2 Alpha and Gamma lineage sequences presented numerous mutations (∼50-80) in comparison to the reference sequence Wuhan-1 (GenBank accession number MN908947.3) (Fig. 5B and C). Analysis of the WTD SARS-CoV-2 Delta lineage sequences revealed similar genetic divergence from the human Delta sequences, however, the number of mutations accumulated in sequences of this lineage in WTD was lower (ranged 40-65) than the observed mutations in the Alpha and Gamma sequences (Fig. 5D). The substitution rate in human SARS-CoV-2 sequences was estimated at 24.4 nucleotide substitutions per year, whereas in WTD the substitution rates were estimated at 35.3, 36.3 and 26.9 substitutions per year for the Alpha, Gamma and Delta sequences, respectively (Fig. S1).

Potential host-adaptive mutations were observed in the Alpha, Gamma and Delta lineages detected in WTD (Tables S4-7). These are non-defining mutations for their respective lineage or sub-lineage and present low frequency among human derived SARS-CoV-2 sequences (global frequency < 0.5%). Mutations in the spike (S) gene are of particular interest due to their potential to result in evasion of immune response. Among the mutations observed in our dataset, S:Q564L was detected in 11 out of 27 (40.7 %) Gamma sequences. This mutation is present in only 167 sequences from the complete GISAID database and in only one sequence out of 68,540 Gamma (as of June 6^th^, 2022). Another mutation in the S gene, S1252F, was detected in nine Alpha (16.4%) and one B.1 lineage sequences from close geographic locations in NY. Among the putative host-adaptive mutations observed in the S gene, four occurred in the receptor-binding domain (RBD): P384S and V445A in one Alpha sequence each, and T323I and Y449H in one Gamma sequence each. No mutations in the S RBD were detected in Delta sequences. Only the RBD V445A mutation, detected in three geographically related Alpha lineage sequences, occurred in more than one sample (6.3%) (Table 4). Other potential host-adaptive mutations were observed in WTD from different regions of NY or in samples from NY and other states (Table S4). For example, the S mutation W258C was detected in eight Alpha variant sequences (27.8%) from WTD from different locations of NY, while S mutation L1203F was detected in four Alpha and three Delta sequences from NY and one B.1.2 from Iowa (IA). Mutations outside the S gene were also observed (Table S5). The ORF1a mutation L4111F is a strong candidate for host-specific adaptation. It was detected in fourteen sequences from NY, including nine Gamma (33.3%) and six Delta sequences, and in five B.1 sequences from WTD from Canada, and one Alpha sequence obtained from WTD in Pennsylvania.

A homoplasy analysis was performed to identify additional sites with potential strong host adaptive mutations of the SARS-CoV-2 VOCs in WTD (Table S6). The analysis identified three mutations (ORF1a:T3150I, S:P25S, and S:T29I) with a consistency index < 0.5, that presented a low frequency in human derived sequences. Other mutations were also detected in WTD from other states and in different lineages (Fig. 6). The mutation in ORF1a:L1853F, was detected in one B.1 and five Alpha sequences from NY, three B.1.311 and 10 B.1.2 sequences from IA and one Delta sequence from Maine (ME) and Canada (Fig. 6A). The ORF1a:L4111F mutation was detected in nine Gamma and one Delta sequences in NY, one Alpha sequence from Pennsylvania (PA), and five B.1 sequences from Canada (Fig. 6B) This mutation is also among mutations above 30% frequency in its respective lineage (Table S5). The S mutation T29I was detected in six Alpha lineage sequences from NY State, one B.1.2 from IA and one Delta sequence from Kansas (KS) (Fig. 6C). Other homoplastic mutations were also detected here mostly on ORF1a and represented by the substitution of Leucine or Serine to Phenylalanine (L or S to F) (Table S6).

**FIG 6.**
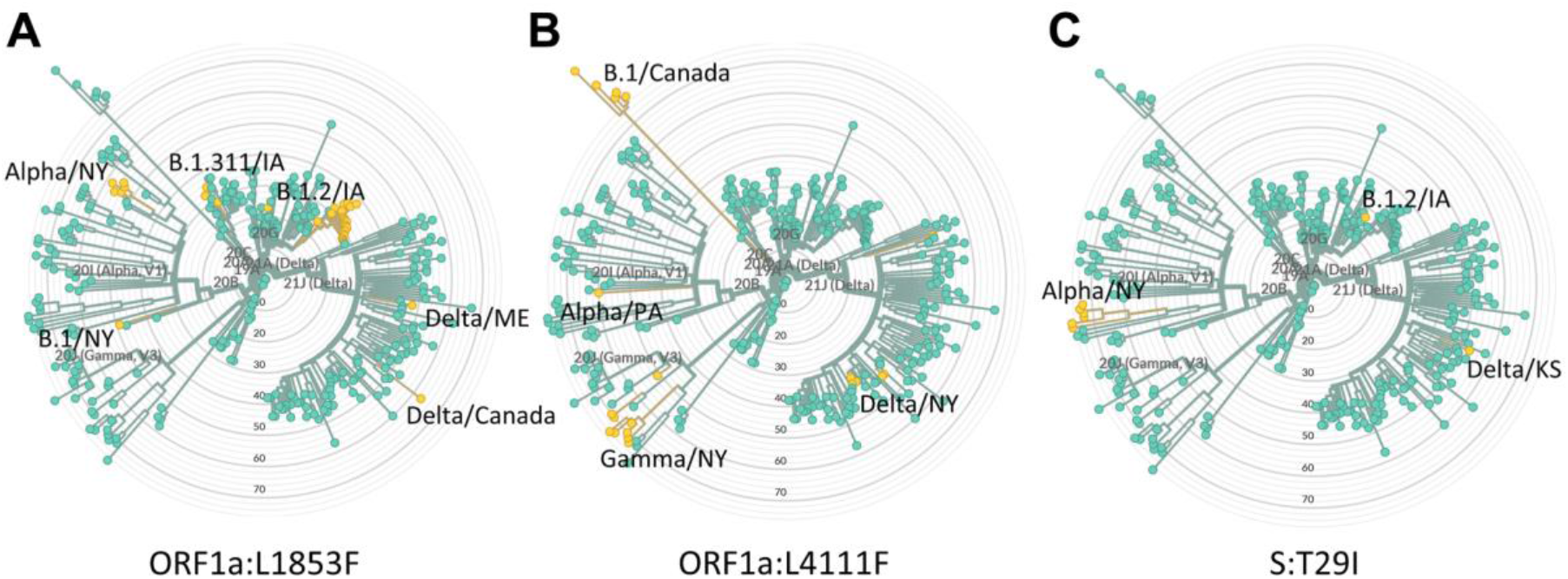
Potential host-adaptive mutations detected in WTD SARS-CoV-2 samples. Mutations highlighted were selected by homoplasy consistency index, prevalence in white-tailed deer derived sequences and detection in multiple geographical regions. Yellow nodes represent sequences with the mutation and green nodes represent the original amino acid. The subclades or nodes where the potential host-adaptive mutations were detected are identified with labels describing the lineage and the state where the samples were collected. (A) Mutation ORF1a:L1853F, (B) mutation ORF1a:L4111F and (C) mutation S:T29I.

### Dispersal of SARS-CoV-2 variants within hotspots of infection in white-tailed deer in New York

The geographic dispersal dynamics of SARS-CoV-2 in high-risk clusters identified using spatial analysis were investigated. The SARS-CoV-2 sequences obtained from seven high risk clusters (C1, C2, C7, C8, C12, C13 and C18) were included in our phylogeographic reconstructions (Fig. 7A). Clusters 7 (n = 10) and 18 (n = 5) comprised a single SARS-CoV-2 variant which were identified as Alpha and Delta, respectively. Cluster 1 (n = 10) contained sequences identified as Alpha (n = 4), Delta (n = 4), and B.1.409 (n = 1) variants. Cluster C2 contained mostly Gamma (n = 27) and one Alpha sequence, while C12 comprised Alpha (n = 7) and Delta variant (n = 6) sequences and C13 comprised Alpha (n = 4), Delta (n = 2) and a single B.1.1 (n = 1) variant sequence.

**FIG 7.**
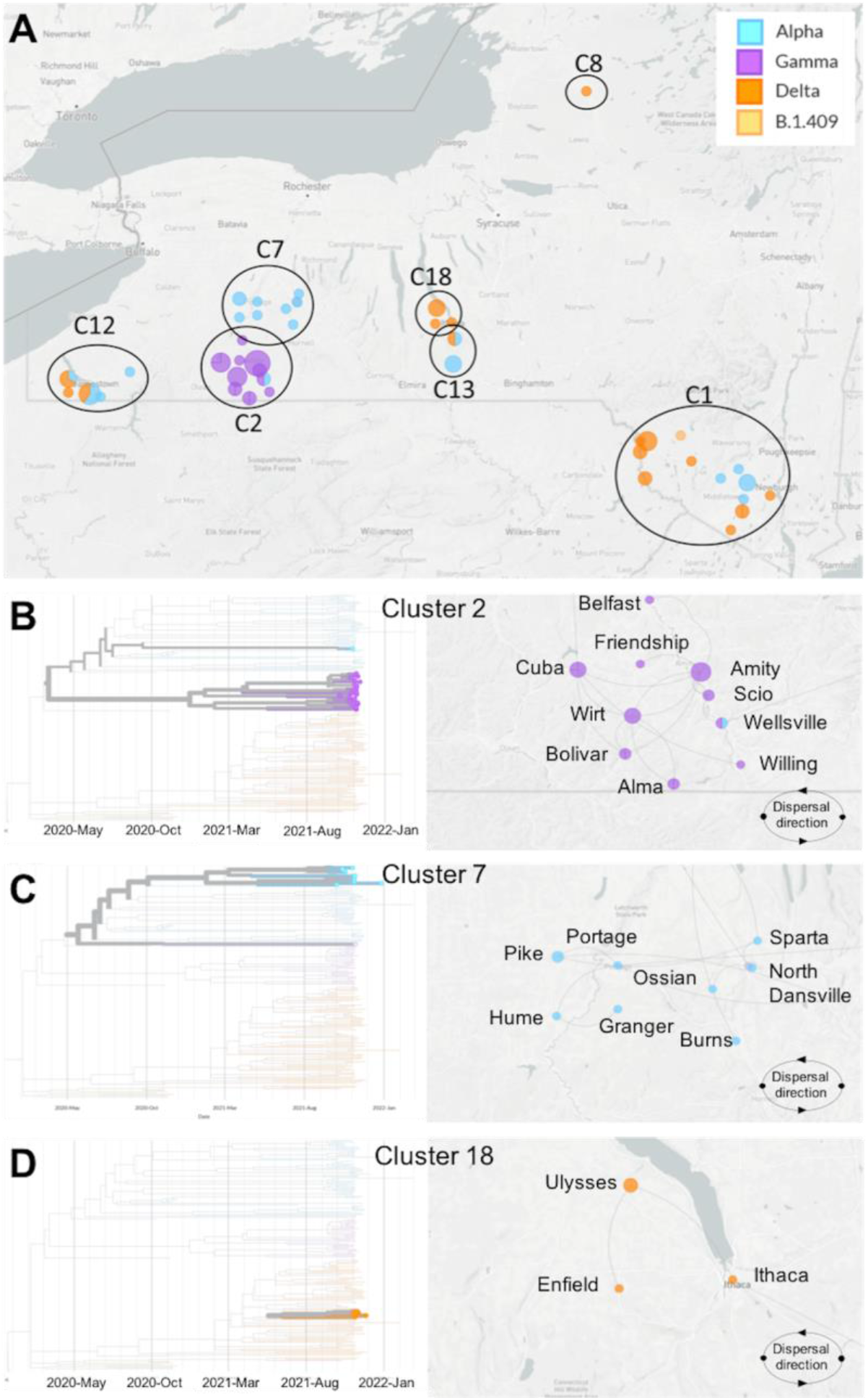
Dispersal of SARS-CoV-2 within hotspots of infection in white-tailed deer in the State of New York (NY). (A) Sampling location and lineage classification of sequences obtained within the seven high-risk clusters in NY. Clusters are identified as C1, C2, C7, C8, C12, C13 and C18. Phylogenetic reconstruction and dispersal analysis of (B) cluster C2, (C) cluster C7 and (D) cluster C18. Directions of dispersal lines are counter-clockwise.

For the phylogeographic dispersal analysis, we focused on C2, C7 and C18 as representative clusters of the three major SARS-CoV-2 lineages identified in our study, Gamma, Alpha and Delta, respectively. The phylogenetic relationship and dispersal pathways were inferred based on whole genome sequences, and the town and date of sample collection. This approach allowed spatial reconstruction of the dispersal history of the viral lineage(s) within each identified cluster. Cluster 2 was geographically restricted to Allegany County and comprised all the Gamma variant sequences detected in our study. These sequences formed a monophyletic branch of closely related sequences (Fig. 7B). Phylogeographic dispersal analysis of the Gamma sub lineages detected in this cluster revealed an intricate dispersal pathway with clear connections between the sequences. Analysis of the directionality of dispersion of the Gamma sequences in C2, point to the samples detected in the town of Cuba as the central focus from which the virus may have spread to Amity, Wirt, and Bolivar and then to other herds in in surrounding towns in the area including Friendship, Willing, and Alma (Fig. 7B).

The Alpha variant C7 involved the counties of Livingston, Wyoming, and Allegany. A total of 10 Alpha variant sequences were recovered from samples within this cluster, which were collected in nine different towns in the region (Dansville, North Dansville, Ossian, Portage, and Sparta in Livingston County; Burns, Hume and Granger in Allegany County; and Pike in Wyoming County). These sequences formed a large branch in the phylogenetic tree that formed two smaller branches of closely related sequences (Fig. 7C). Dispersal analysis of the Alpha sequences detected in this cluster revealed two potential dispersal pathway(s) with links between the detected sequences. It appears that the sample detected in Portage may be the central foci in C7 from where the virus may have spread to eastern (Sparta, North Dansville and Ossian) and western locations (Pike, Hume and Granger) (Fig. 7C).

The Delta variant C18 represents the smallest (n = 5) of the clusters in which we analyzed the spatial dispersal of SARS-CoV-2. This cluster was restricted to Tompkins County and samples were detected in the towns of Ulysses, Ithaca, and Enfield. The Delta sequences detected in C18 formed a unique branch in the phylogenetic tree with highly related sequences (Fig. 7D). Phylogeographic dispersal analysis of the Delta sequences detected in this cluster revealed a clear connection between the sequences with the samples detected in the town of Ulysses appearing to be the central foci of the cluster which further dispersed into nearby Enfield and Ithaca (Fig. 7D). The phylogenetic relationship of the SARS-CoV-2 sequences detected in WTD and their divergence from contemporary samples recovered from humans, demonstrate the circulation and indicate onward transmission of Alpha, Gamma, and Delta SARS-CoV-2 VOCs in WTD in NY.

## DISCUSSION

White-tailed deer are highly susceptible to SARS-CoV-2 infection and efficiently transmit the virus to direct or indirect contact animals as demonstrated by our research group and collaborators from the National Animal Disease Center (NADC) (8, 10). Follow up testing for SARS-CoV-2 in captive and free-ranging WTD in different regions of the USA and Canada have shown that a significant proportion of WTD may have been exposed or infected with SARS-CoV-2 in North America (12, 14, 15, 17).

In the present study, we used samples from a well-established surveillance program for CWD in wild deer in the State of NY (20), to conduct a comprehensive investigation and assess the prevalence of SARS-CoV-2 infection in WTD over two deer hunting seasons (2020 and 2021) during the COVID-19 pandemic. During these seasons 253,990 and 211,269 WTD were harvested by hunters in NY, respectively with ∼1% of the animals being sampled and tested for CWD (2,700 and 2,762, respectively) and subsequently included in our study. Of the samples tested for SARS-CoV-2 17 (0.6%) were positive in 2020 and 583 (21.1%) in 2021. The sampling strategy used by the NYS Department of Environmental Conservation (NYSDEC) emphasizes locations and age classes of deer that are at greater risk for CWD infection using weighted surveillance and a qualitative risk assessment. The sampling calculations are determined by three primary criteria: i. deer sex and age classes, with emphasis given to older bucks and does; ii. proximity to geographic risk factors, such as captive cervid facilities, taxidermy and deer processing centers, and proximity with CWD-endemic states (like PA); and finally, iii. the deer population size, with areas with larger deer counts (estimated based on the number of deer harvested by square kilometer) receiving more intense surveillance. This sampling strategy allowed us to evaluate the infection rate and dynamics of SARS-CoV-2 infection using samples that are representative of the adult WTD population distributed across the State of NY (∼1M WTD). Our results showed a striking 35-fold increase in SARS-CoV-2 prevalence and consequently a broader geographic distribution of the infection between 2020 (10/62 positive counties, Season 1) and 2021 (57/62 positive counties, Season 2).

Spatial clustering analysis based on positive SARS-CoV-2 detections revealed several hotspots or areas with a high concentration of cases in WTD in NYS. Notably, when the viral sequence information was compared with the spatial clusters, several clusters of closely related SARS-CoV-2 variants overlapped with these defined geographic hotspots, indicating local circulation and transmission of the virus within the deer populations in these areas. These findings corroborate observations of early studies which found evidence of potential transmission of SARS-CoV-2 amongst wild WTD in the states of Ohio and Iowa in the US or in the province of Ontario in Canada (12, 15, 21). Interestingly, the broad geographic distribution of the Alpha and Delta SARS-CoV-2 variants in our study and the identification of several clusters of closely related viruses within confined geographic locations, highlight the occurrence of multiple viral spillover events from humans to WTD between 2020 and 2021, which was followed by deer-to-deer transmission of the virus.

Transmission of the virus between WTD can occur through direct contact or indirectly, likely through respiratory droplets (8, 10). While assessing SARS-CoV-2 transmission pathways within WTD populations, animal age and sex must be considered as direct or indirect contact between animals can be increased by distinct social behaviors of different age groups and sex. For example, maternal groups consisting of females and fawns are relatively isolated during the spring and summer while males typically have large home ranges and have increased movement and contact with other deer especially during the breeding season (16, 22). Interestingly, results from our study, in which sample collection overlaps with the breeding season (September to December), indicate that adult male deer are at significantly higher risk of becoming infected with SARS-CoV-2 than other sex and age classes. These observation are consistent with a previous serological study that has shown higher prevalence of antibodies in adult male WTD (16) and highlight the potential epidemiological role of males in maintenance and transmission of the virus within wild WTD populations. Although the pathways for transmission of SARS-CoV-2 from humans-to-WTD remain largely unknown, human activities such as feeding wildlife or targeted baiting of hunting prey (e.g., WTD) could provide the opportunity for human-to-WTD transmission of the virus. Animal feed and feed ingredients are known to promote and enhance survival of several animal viruses, including coronaviruses (23), thus wildlife feeding practices should be investigated as a risk factor and avoided due to their potential to enable spillover of SARS-CoV-2 from humans to WTD and perhaps other susceptible wildlife species (e.g. mink, deer mice, raccoon dogs, etc.; Fig. 8) (24–26).

**FIG 8.**
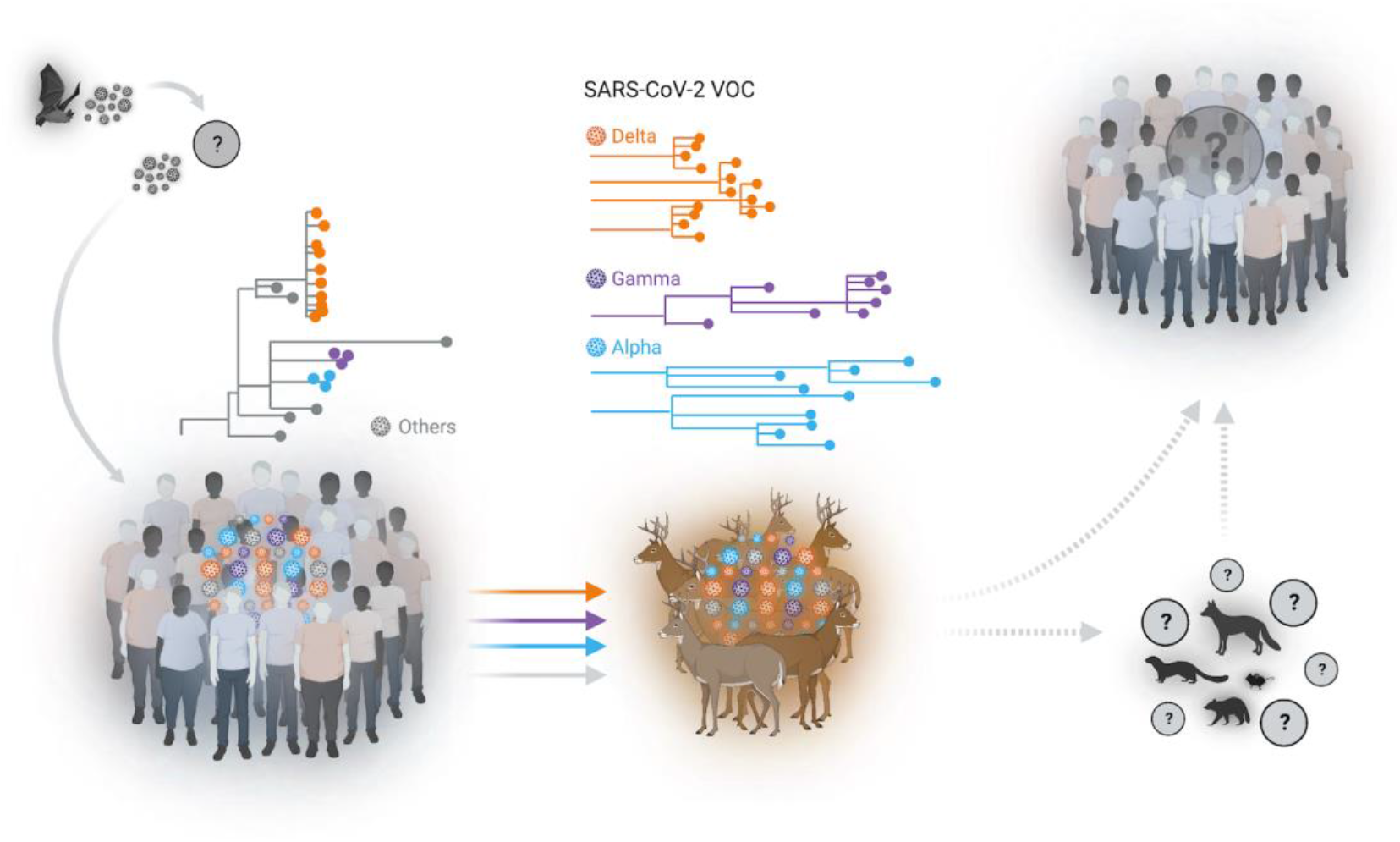
Spillover of SARS-CoV-2 from humans to white-tailed deer. Multiples spillover events of SARS-CoV-2 from humans to wild white-tailed deer (WTD) population were identified in several geographic areas in the State of New York (NY). Sequence analysis of SARS-CoV-2 genomes demonstrated the circulation of classical SARS-CoV-2 lineages as well as the co-circulation of three major variants of concern (VOCs) (Alpha, Gamma, and Delta) in WTD. Our analysis suggests the occurrence of multiple spillover events (human-to-deer) of multiples SARS-CoV-2 lineages with subsequent deer-to-deer transmission and emergence of highly genetically diverse viruses in this species. The potential for spillback of variant SARS-CoV-2 to humans or yet spillover from WTD to other susceptible wildlife species wildlife species (e.g. mink, deer mice, raccoon dogs, and red fox) remains unknown.

In our study, we have not found direct sequence links between the viruses circulating in WTD in NY with any of the available SARS-CoV-2 sequences recovered from humans. In fact, the viral sequences obtained from WTD are highly divergent from available human-derived SARS-CoV-2 sequences, with some strains accumulating over 70 nucleotide mutations across the genome. High genetic divergence from human SARS-CoV-2 sequences was observed amongst sequences from all three major (Alpha, Gamma and Delta) variants identified in our study. Notably, Alpha and Gamma sequences presented the highest divergence to human sequences, which aligns with the fact that these variants were likely introduced earlier and circulated longer in WTD populations than Delta variant viruses. The peak of detection of Alpha and Gamma variants in humans in NY occurred between April and June 2021, with only sporadic detections occurring after August, while the Delta variant was the dominant variant circulating in humans in NY during the time of sample collection from WTD included in our study. These findings indicate independent evolution of SARS-CoV-2 VOCs in WTD and demonstrate parallel circulation of viral variants (Alpha and Gamma) that had been replaced by the Delta variant in humans, thus highlighting the potential that WTD may serve as a wildlife reservoir for nearly extinct SARS-CoV-2 variants. Thus, it is extremely important to continue to monitor WTD populations in the areas sampled in our study to assess the dynamics of the identified VOCs as well as of other deer-associated variants that might evolve and emerge in this host and potentially spillback into humans.

To date, only one report described the detection of a single WTD-like SARS-CoV-2 variant recovered from humans in Canada, which suggests potential deer-to-human transmission of the virus (14). Deer hunting is a widespread practice in North America that is regulated by the states and used as a method of population control for this highly prolific animal species. In 2020, for example ∼6.5 million WTD were harvested in the US and Canada (27). This practice provides opportunities for close contact between a potentially infected reservoir and humans, along with wildlife rehabilitation, captive cervid ownership, and zoological collections. In the present study, we recovered infectious virus from seven RPLN samples that tested positive for SARS-CoV-2 RNA, corroborating findings from previous studies that detected infectious virus in samples collected from wild WTD (12, 15). Although the proportion of live virus positive samples detected was low, the findings are important in that they demonstrate that there is a risk of contact with infectious SARS-CoV-2 during handling and processing of WTD carcasses, which could lead to spillback via deer-to-human transmission of the virus. Our previous experimental data, assessing the infectious and transmission dynamics of SARS-CoV-2 in WTD, demonstrated a short window (∼5 days) of infectiousness, a period in which experimentally inoculated animals excrete the virus in oral and nasal secretions and carry infectious/replicating virus in respiratory and lymphoid tissues and are able to transmit the virus to contact animals (10). The short viral replication window in tissues may have contributed to the low detection of infectious virus in the samples tested in our study. Additional factors such as a longer time from sample collection to testing and sub-optimal sample storage conditions may also have negatively affected the success of virus isolation in our study.

The high number of mutations observed in SARS-CoV-2 samples recovered from WTD in relation to human samples indicate that the virus has been circulating and evolving within the deer population. Some of the observed mutations could be a response to host adaptation. The mutation ORF1a:L4111F, for example, is observed in WTD-derived samples from multiple geographic regions and is present at a low frequency (< 0.5%) among all human sequences available in GISAID database. This suggests that this mutation emerged independently in deer after spillover from humans. The absence or very low frequency of fixed mutations in the S gene suggests that host-specific adaptation was not necessary for human-to-deer spillover, reinforcing the notion of SARS-CoV-2 as a “generalist” virus (28). The presence of homoplasious and high frequency mutations in ORF1a and ORF1b emphasize the importance of also characterizing non-Spike protein mutations, as they may also play a role on virus host-range and species specificity. A few mutations detected in the S gene lie near aa residues that define the Delta, Alpha and Gamma VOCs (Table S7). The mutation S:R683W detected in Delta WTD sequences, for example, is located close to the S:P681R site, which enhances cleavage of the S into the S1 and S2 subunits (29). Two other substitutions detected in Alpha WTD samples, S:D568N and S:I569T, lie in close proximity to the S:A570D mutation, which may play an important role modulating the conformation of S RBD, and was shown to contribute to Alpha VOC infectivity in humans (30). It is possible that these S substitutions might increase affinity of S protein for the WTD ACE2 receptor orthologue (31), however additional experiments are needed to determine their actual biological function and whether they contribute to SARS-CoV-2 infection, replication and transmission.

The evidence generated in this study demonstrate widespread dissemination of SARS-CoV-2 in wild WTD populations across the State of NY, and further indicate the co-circulation of three major VOCs in this new wildlife host. Notably, circulation of SARS-CoV-2 in WTD results in emergence of highly genetically diverse viruses in this species. These observations highlight the need to establish continuous surveillance programs to monitor the circulation, distribution, and evolution of SARS-CoV-2 in WTD populations and to stablish measures to minimize additional virus introductions in animals that may lead to spillback of novel deer-adapted SARS-CoV-2 variants to humans.

## MATERIAL AND METHODS

### Sample collection

Retropharyngeal lymph nodes (RPLN) from white-tailed deer (WTD) from the Chronic Wasting Disease (CWD) surveillance program in New York State (NYS) were used in this study to investigate the presence of SARS-CoV-2. Samples were collected and submitted to Cornell Animal Health Diagnostic Center (AHDC) for ELISA testing. A total of 5,462 RPLN collected during the hunting seasons of 2020 (Season 1, September - December 2020, n = 2,700) and 2021 (Season 2, September - December 2021, n = 2,762) were included in this study. The WTD sex, age, date of harvest, and location to the level of town were collected with the sample to use for analysis.

### Nucleic acid extraction and rRT-PCR

RPLN tissues (200 mg) were minced with a sterile scalpel, resuspended (1/10 w/v) in PBS and homogenized using magnetic beads. For the RNA extraction, 200 µl of the homogenate was used as input using the MagMax Core extraction kit (Thermo Fisher, Waltham, MA, USA) and the automated KingFisher Flex nucleic acid extractor (Thermo Fisher, Waltham, MA, USA) following the manufacturer’s recommendations. The real time reverse transcriptase PCR (rRT-PCR) was performed using the EZ-SARS-CoV-2 real-time RT-PCR assay (Tetracore Inc., Rockville, MD), which detects both genomic and sub-genomic viral RNA for increased diagnostic sensitivity. An internal inhibition control was included in all reactions. Positive and negative amplification controls were run side by side with test samples.

### Virus isolation

For virus isolation, RPLN that tested positive for SARS-CoV-2 by rRT-PCR with Ct values lower than 30 were subjected to virus isolation under biosafety level 3 conditions at Cornell University. Vero E6/TMPRSS2 (JCRB Cell Bank, JCRB1819) were cultured in Dulbecco’s modified eagle medium (DMEM) supplemented with 10% fetal bovine serum (FBS), L-glutamine (2mM), penicillin (100 U.ml^−1^), streptomycin (100 μg.ml^−1^) and geneticin (1 mg.ml^- 1^). Twenty-four well plates were seeded with ∼75,000 Vero E6/TMPRSS2 cells per well 24 h prior to sample inoculation. Cells were rinsed with phosphate buffered saline (PBS) (Corning) and inoculated with 200 μl of the homogenate and inoculum adsorbed for 1 h at 37 °C with 5% CO_2_. Mock-inoculated cells were used as negative controls. After adsorption, replacement cell culture media was added, and cells were incubated at 37 °C with 5% CO_2_ and monitored daily for cytopathic effect (CPE) for 3 days. SARS-CoV-2 infection in CPE-positive cultures was confirmed with an immunofluorescence assay (IFA) as described previously (8). Cell cultures with no CPE were frozen, thawed, and subjected to two additional blind passages/inoculations in Vero E6/TMPRSS2 cell cultures. At the end of the third passage, the cells cultures were subjected to IFA.

### Data acquisition

All WTD derived SARS-CoV-2 genomes (n = 159) were retrieved from GISAID (accessed May 1, 2022). All human derived SARS-CoV-2 genomes (n = 91,796) from the State of NY, US were retrieved, and a local BLAST database was built to obtain 3837 sequences with highest nucleotide similarities to WTD derived SARS-CoV-2 genomes in current study.

### SARS-CoV-2 genome sequencing

Total RNA was reverse transcribed using SuperScript IV Reverse Transcriptase (ThermoFisher scientific) as described in the protocol available at dx.doi.org/10.17504/protocols.io.br54m88w. The cDNA from each sample was used four times for parallel generation of tiled amplicons with ARTIC V3 and ARTIC V4.1 primers (IDT) and library preparation using a modified ARTIC network’s nCoV-2019 sequencing protocol v2 (32). Minor modifications include the use of Q5 High-Fidelity 2X Master Mix and the use of 4µl cDNA in a 25µl PCR reaction. Additionally, the annealing and denaturation temperatures were dropped to 64 °C and 95 °C, respectively. Purified amplicons were diluted 1:10 and quantified before used as input for two sequencing library preparations (V3 and V4.1 primers), that were loaded on R9.4 flow cells in a MinION Mk1B (Oxford Nanopore Technologies, ONT).

### Genomic analysis

Raw FAST5 reads were basecalled and demultiplexed using Guppy v5.0.16 in high accuracy mode. The output FASTQ files were processed through the ARTIC bioinformatics pipeline using the applicable primer scheme (V3 or V4.1) to generate consensus sequences (https://github.com/artic-network/artic-ncov2019). Lineage classification was performed using Pangolin version 4.0.6 (19). Only sequences passing Pangolin QC thresholds for minimum length and maximum N content were used for downstream analyses.

### Interactive phylogenomic and phylogeographic analyses

For phylogenetic analysis of WTD derived SARS-CoV-2 genomes, two curated datasets were created. First dataset consisted of all 164 SARS-CoV-2 from current study and 159 deer derived SARS-CoV-2 genomes downloaded from GISAID database. The second dataset consisted of 3837 human derived SARS-CoV-2 genomes and 323 deer derived SARS-CoV-2 genomes from the present study and GISAID database. Phylogenetic analysis on both datasets was performed by using procedures implemented in Nextstrain (33). Briefly, multiple sequence alignment was performed using Nextalign, maximum likelihood tree inferred using IQ-TREE through Augur toolkit, and data visualization through Auspice. Maps in figures 3A and 3B were generated using Microreact (34).

### Mutation analysis

Sequences from the present study were grouped according to its VOC classification and nucleotide sequence alignments were performed individually for each group of VOCs using MAFFT v7.453 (35). Variations in relation to reference genome (GenBank accession number MN908947.3) were identified in Geneious Prime 2019 software. Lineage-defining non-synonymous mutations were obtained from the Lineage Comparison tool of Outbreak web interface (36) and excluded from the dataset. The remaining mutations were used to screen for possible host-adaptive mutations. The phylogenetic tree with WTD derived sequences was screened for homoplasies using HomoplasyFinder to strengthen findings of putative host-adaptive mutations (37). Nucleotide positions with a consistency index <0.5 were considered for further analysis.

### Time scale distribution of human derived SARS-CoV2

To place the deer derived SARS-CoV-2 VOCs from current study in the context of VOCs circulating in human population, the epidemiological metadata (VOCs and date of collection) of a total of 91,000 (NY) and 250,000 (USA) human derived SARS-CoV-2 genome sequences from February 2020 to May 2022 were retrieved from GISAID. A stacked bar graph showing monthly cases of different VOCs in human population of NY and USA was created (34) Using GraphPad prism 9 (GraphPad Software, San Diego, CA, USA)

### SaTScan spatial analysis Relative Risk and Logistic Regression Analysis

The space-time discrete Bernoulli model (STBM) within SaTScan software version 10.0.2 (http://www.satscan.org) (38) was used to identify statistically significant clusters of SARS-CoV-2 infection in WTD population. For each sample, the geographic location (latitude and longitude) was determined based on the centroid of town using the US Census Bureau County TIGER/Line Shapefile for County Subdivisions, which are equivalent to towns, cities and other similar administrative divisions. To detect high rates of SARS-CoV-2 in deer clusters, the Monte Carlo hypotheses test method and 999 replications were applied to detect a high prevalence rate of SARS-CoV-2 (39). The spatial clusters were visualized in QGIS (geographic information system) mapping software version 3.16.16 (40).

### Relative Risk and Logistic Regression Analysis

The measure of relative risk (RR) was used to identify locations with a greater risk of having SARS-CoV-2 in deer, following the equation RR=(c/e)/((C-e)/(C-e)) where c represents the total number of observed cases in the town, e represents the total number of expected cases in a town, C is the total number of observed cases in New York State. If the RR was larger than 1, the location had a greater risk of having a positive in deer. A logistic regression analysis conducted on deer outside the clusters was used to determine whether the difference between deer grouped by age, sex, and season makes sense in the comparison of age, sex, and season for SARS-CoV-2 infection.

## Data availability

All SARS-CoV-2 consensus genomes are deposited in GISAID, https://www.gisaid.org/; Accession numbers are available in Table S8 and raw reads have been submitted to NCBI’s Short Read Archive (BioProject Number PRJNA872140).

## Acknowledgments

We thank the Cornell Biosafety team for the support. We also thank the Animal Health Diagnostic Center at Cornell University for the use of extraction and real-time PCR equipment. The NYS Department of Environmental Conservation was instrumental in tissue collection and cooperation in data sharing for this study. Tetracore Inc. generously provided the real-time PCR reagents used to test the samples in this study.

