## Supplemental Figure 1 for "White-tailed deer (*Odocoileus virginianus*) may serve as a wildlife reservoir for nearly extinct SARS-CoV-2 variants of concern"

### Supplementary figure 1

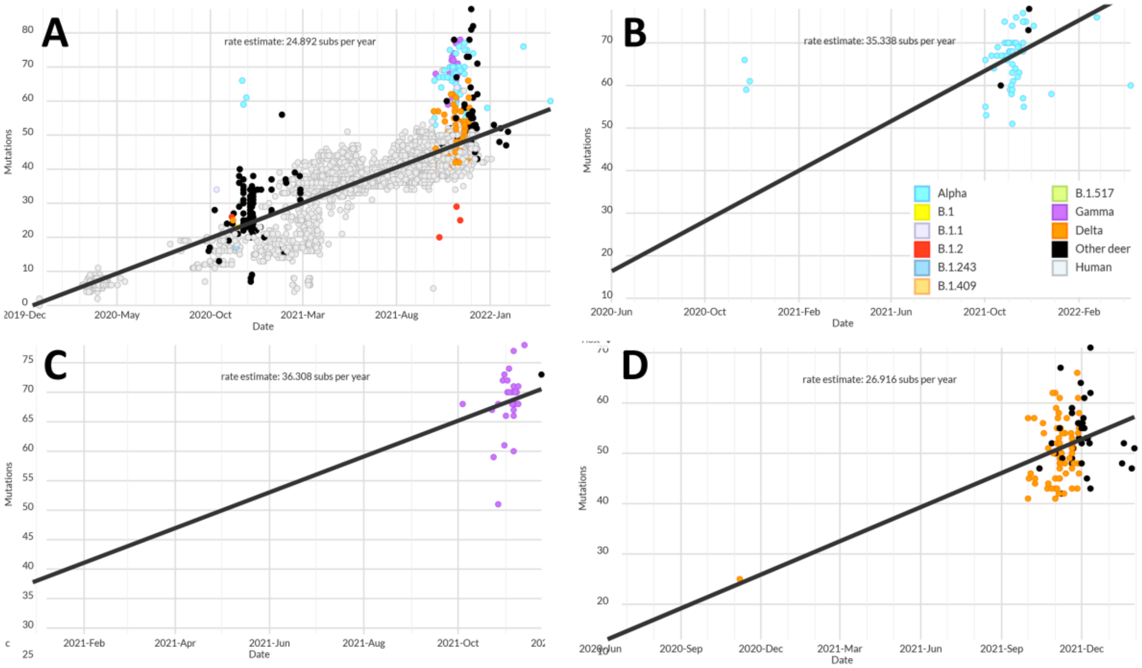


**SUPP FIG 1** Rate estimates of substitutions per year. Colored dots (sky blue: Alpha; purple: Gamma and orange: Delta) represent SARS-CoV2 variants of concern (VOCs) derived from WTD from this study, black dots represent the Pan-US SARS-CoV-2 sequences derived from white-tailed deer (WTD) and grey dots represent SARS-CoV-2 sequences derived from human. The Y axis shows the number of mutations compared to the reference sequence Wuhan-1 (GenBank accession number MN908947.3). (A) Substitutions rate of all deer-derived sequences and 3837 human-derived sequences, (B) Alpha VOC substitution rate in deer, (C) Gamma VOC substitution rate in deer and (D) Delta VOC substitution rate in deer.
