## Supplemental Tables for "White-tailed deer (*Odocoileus virginianus*) may serve as a wildlife reservoir for nearly extinct SARS-CoV-2 variants of concern"

**S Table 1.** Number of retropharyngeal lymph nodes from white-tailed deer tested in each county in the State of New York (NY) in Seasons 1 (September - December 2020, n = 2700) and Seasons 2 (September - December 2021, n = 2762).

| **County** | **Tested 2020** | **Positive 2020** | **Tested 2021** | **Positive 2021** |
| --- | --- | --- | --- | --- |
| Albany | 19 | 0 | 12 | 4 |
| Allegany | 157 | 0 | 148 | 70 |
| Broome | 98 | 0 | 97 | 14 |
| Cattaraugus | 115 | 0 | 143 | 32 |
| Cayuga | 48 | 0 | 42 | 11 |
| Chautauqua | 123 | 0 | 228 | 62 |
| Chemung | 75 | 1 | 82 | 16 |
| Chenango | 56 | 0 | 52 | 12 |
| Clinton | 69 | 0 | 56 | 1 |
| Columbia | 33 | 0 | 33 | 4 |
| Cortland | 27 | 0 | 21 | 1 |
| Delaware | 77 | 1 | 78 | 0 |
| Dutchess | 43 | 0 | 31 | 5 |
| Erie | 109 | 0 | 84 | 17 |
| Essex | 32 | 0 | 40 | 7 |
| Franklin | 35 | 0 | 24 | 2 |
| Fulton | 7 | 0 | 12 | 0 |
| Genesee | 45 | 0 | 32 | 5 |
| Greene | 23 | 0 | 61 | 2 |
| Hamilton | 7 | 0 | 9 | 1 |
| Herkimer | 28 | 0 | 20 | 0 |
| Jefferson | 42 | 0 | 39 | 8 |
| Lewis | 30 | 0 | 22 | 10 |
| Livingston | 52 | 0 | 69 | 23 |
| Madison | 38 | 0 | 39 | 10 |
| Monroe | 41 | 2 | 43 | 9 |
| Montgomery | 16 | 0 | 10 | 0 |
| Nassau |  |  | 5 | 0 |
| Niagara | 65 | 0 | 39 | 4 |
| Oneida | 65 | 0 | 55 | 6 |
| Onondaga | 55 | 0 | 49 | 9 |
| Ontario | 85 | 0 | 40 | 7 |
| Orange | 66 | 0 | 70 | 32 |
| Orleans | 32 | 1 | 31 | 6 |
| Oswego | 64 | 0 | 46 | 2 |
| Otsego | 29 | 0 | 35 | 8 |
| Putnam | 13 | 0 | 17 | 0 |
| Rensselaer | 24 | 0 | 53 | 12 |
| Rockland | 5 | 0 | 27 | 2 |
| Saratoga | 16 | 0 | 24 | 0 |
| Schenectady | 10 | 0 | 7 | 0 |
| Schoharie | 26 | 0 | 29 | 3 |
| Schuyler | 21 | 0 | 22 | 3 |
| Seneca | 50 | 0 | 29 | 1 |
| St. Lawrence | 49 | 0 | 37 | 5 |
| Steuben | 120 | 4 | 143 | 30 |
| Suffolk | 24 | 0 | 23 | 1 |
| Sullivan | 64 | 2 | 97 | 45 |
| Tioga | 44 | 0 | 60 | 19 |
| Tompkins | 29 | 0 | 33 | 12 |
| Ulster | 41 | 1 | 47 | 14 |
| Warren | 13 | 0 | 13 | 0 |
| Washington | 40 | 0 | 56 | 4 |
| Wayne | 52 | 0 | 52 | 10 |
| Westchester | 24 | 0 | 58 | 1 |
| Wyoming | 59 | 0 | 55 | 15 |
| Yates | 24 | 0 | 22 | 1 |

**S Table 2.** The presence of SARS-CoV-2 in white-tailed deer in the State of New York has been observed in spatial clusters in the hunting Season 2 (September - December 2021)^1^.

| cluster | Total IUMPS^2^ | Radius (Km) | Observed  (O) | Expected  (E) | O/E | Relative  Risk (RR) | Log-Likelihood  Ratio | p-Value^3^ |
| --- | --- | --- | --- | --- | --- | --- | --- | --- |
| C1 | 29 | 55.31 | 89 | 36.86 | 2.41 | 2.68 | 41.029 | <0.001 |
| C2 | 14 | 21.70 | 56 | 22.12 | 2.53 | 2.70 | 27.62 | <0.001 |
| C3 | 57 | 67.98 | 0 | 20.85 | 0 | 0 | 23.91 | <0.001 |
| C4 | 22 | 34.75 | 0 | 19.59 | 0 | 0 | 22.43 | <0.001 |
| C5 | 26 | 80.43 | 1 | 19.80 | 0.049 | 1.1 | 18.42 | <0.001 |
| C6 | 20 | 64.38 | 0 | 15.38 | 0 | 0 | 17.53 | <0.001 |
| C7 | 17 | 21.17 | 30 | 11.79 | 2.54 | 2.63 | 14.58 | <0.001 |
| C8 | 6 | 17.89 | 14 | 4.21 | 3.32 | 3.38 | 11.11 | 0.016 |
| C9 | 12 | 25.18 | 0 | 9.27 | 0 | 0 | 10.50 | 0.040 |
| C10 | 24 | 36.33 | 1 | 10.95 | 0.091 | 0.090 | 8.79 | 0.154 |
| C11 | 20 | 28.30 | 2 | 13.06 | 0.15 | 0.15 | 8.60 | 0.164 |
| C12 | 17 | 27.21 | 50 | 29.49 | 1.70 | 1.76 | 8.42 | 0.187 |
| C13 | 7 | 16.99 | 13 | 4.63 | 2.81 | 2.85 | 7.54 | 0.404 |
| C14 | 12 | 22.57 | 7 | 6.11 | 0 | 0 | 6.90 | 0.585 |
| C15 | 8 | 19.84 | 0 | 5.05 | 0 | 0 | 5.70 | 0.921 |
| C16 | 12 | 29.41 | 0 | 4.63 | 0 | 0 | 5.22 | 0.983 |
| C17 | 10 | 20.02 | 1 | 6.5 | 0.14 | 0.14 | 4.68 | 0.996 |
| C18 | 3 | 10.66 | 8 | 2.95 | 2.71 | 2.74 | 4.83 | 0.999 |
| C19 | 13 | 46.33 | 1 | 6.53 | 0.15 | 0.15 | 4.26 | 0.999 |

^1^ Results of the spatial clusters model

^2^ IUMPS (Internal database ID for unique towns from the IPUMS geospatial dataset, which is based on the US Census TIGER data)

^3^ Statistically significant at p≤0.005.

**S Table 3.** Results of the logistic regression model using demographic and seasonal factors for SARS-CoV-2 testing in white-tailed deer positive in the State of New York, Season 1 (September - December 2020) and Season 2 (September - December 2021)^1^.

| Response Variable | Predictor variables | OR^2^ | 95% CI^3^ | P-Value^4^ |
| --- | --- | --- | --- | --- |
| Testing Result | Season 2021-22  Yearling  Male | 59.04  0.865  1.952 | 34.83-111.08  0.70-1.059  1.591-2.407 | <0.01  0.164  <0.01 |
| Testing Result | Yearling  Male | 0.876  1.906 | 0.711-1.07  1.55-2.355 | 0.210  <0.01 |

^1^ Outcome (N =	SARS-CoV-2 cases from 5,462 deer samples)

^2^ OR- Odds ratio

^3^ CI- Confidence Interval of the OR

^4^ Statistically significant at P≤0.05

**S Table 4.** Mutations above 4% frequency in each variant of concern (VOC) detected in the spike gene of New York (NY) white-tailed deer (WTD) SARS-CoV-2 with frequencies < 1% in GISAID database.

| Lineage | nt. position | Region | a.a mutation | Proportion in VOC of NY WTD^1^ | Number of GISAID sequences with the mutation^2^ | Proportion in lineage or sublineages |
| --- | --- | --- | --- | --- | --- | --- |
| Alpha | 21604 | S | Q14H | 9.3 | 7258 | <0.5% |
| Delta | 21635 | S | P25S | 6 | 4331 | 1% |
| Alpha | 21648 | S | T29I | 9.3 | 10751 | <0.5% |
| Delta | 21806 | S | P82T | 6.3 | 1021 | 1% |
| Alpha | 21853 | S | K97N | 8.3 | 2995 | <0.5% |
| Alpha | 21969 | S | C136S | 11.4 | 50 | <0.5% |
| Alpha | 22095 | S | D178G | 6.7 | 1095 | <0.5% |
| Alpha | 22115 | S | N185D | 10.9 | 1622 | <0.5% |
| Alpha | 22118 | S | F186L | 15.6 | 1159 | <0.5% |
| Alpha | 22336 | S | W258C | 27.8 | 441 | <0.5% |
| Alpha | 22896 | S | V445A | 6.3 | 738 | <0.5% |
| Gamma | 23253 | S | Q564L | 40.7 | 167 | <0.5% |
| Alpha | 23264 | S | D568N | 5.5 | 297 | <0.5% |
| Alpha | 23268 | S | I569T | 5.5 | 109 | <0.5% |
| Delta | 23609 | S | R683W | 5.7 | 1516 | <0.5% |
| Alpha | 24054 | S | A831V | 7.3 | 5776 | <0.5% |
| Gamma | 24086 | S | G842S | 11.1 | 1611 | <0.5% |
| Gamma | 24372 | S | S937L | 7.7 | 37 | <0.5% |
| Alpha | 24914 | S | D1118Y | 12.7 | 6222 | <0.5% |
| Alpha | 25169 | S | L1203F | 7.3 | 2256 | <0.5% |
| Delta | 25169 | S | L1203F | 4.3 | 2256 | 1% |
| Alpha | 25184 | S | Q1208E | 7.3 | 515 | <0.5% |
| Alpha | 25317 | S | S1252F | 16.4 | 9185 | <0.5% |

^1^ Proportion in sequences obtained in the present study

^2^ As of June 6th 2022

**S Table 5.** Mutations above 30% frequency in each variant of concern (VOC) of New York (NY) white-tailed deer (WTD) SARS-CoV-2 with frequencies < 1% in GISAID database.

| Lineage | Nt. position | Region | A.a mutation | Proportion in VOC of NY WTD^1^ | Number of GISAID sequences with the mutation^2^ | Proportion in lineage or sublineages |
| --- | --- | --- | --- | --- | --- | --- |
| Alpha | 2207 | Orf1a | I648L | 27.8 | 2116 | <0.5% |
| Alpha | 28011 | ORF8 | H40Y | 38.5 | 3871 | <0.5% |
| Gamma | 2937 | Orf1a | T891I | 100 | 7882 | 1% |
| Gamma | 6723 | Orf1a | T2153I | 44 | 7390 | <0.5% |
| Gamma | 11511 | Orf1a | T3749I | 33.3 | 1880 | 1% |
| Gamma | 12596 | Orf1a | L4111F | 33.3 | 1374 | <0.5% |
| Gamma | 14575 | Orf1b | A370S | 100 | 1154 | <0.5% |
| Gamma | 17738 | Orf1b | T1424I | 55.6 | 2443 | <0.5% |

^1^ Proportion in sequences obtained in the present study

^2^ As of June 6^th^ 2022

**S Table 6.** Homoplasious mutations detected in this study and other studies, with frequencies detected in New York (NY) white-tailed deer (WTD).

| Lineage | nt position | Region |  | a.a mutation | Proportion in VOC of NY WTD^1^ | | Reference^2^ |
| --- | --- | --- | --- | --- | --- | --- | --- |
| Alpha | 5822 | Orf1a |  | L1853F | 9.1 | (1), (2) | |
| Alpha | 9510 | Orf1a |  | T3082I | 1.9 | (1) | |
| Alpha | 9510 | Orf1a |  | 3082 | 1.9 | (1) | |
| Gamma | 9711 | Orf1a |  | S3149F | 12 | (1) | |
| Delta | 9711 | Orf1a |  | S3149F | 2.9 | (1) | |
| Delta | 9714 | Orf1a |  | T3150I | 1.4 | Present study | |
| Alpha | 11195 | Orf1a |  | L3644F | 14 | (1) | |
| Delta | 11195 | Orf1a |  | L3644F | 1.4 | (1) | |
| Alpha | 15451 | Orf1b |  | G662S | 1.8 | (1) | |
| Gamma | 21618 | S |  | T19I | 34.6 | (1) | |
| Delta | 21635 | S |  | P25S | 6 | Present study | |
| Alpha | 21648 | S |  | T29I | 9.3 | Present study | |
| Delta | 25563 | ORF3a |  | Q57H | 1.4 | (1) | |

^1^ Proportion within VOC in sequences obtained in the present study

^2^ Study that identified mutation as homoplasious

**S Table 7.** Detection of mutations close to variant of concern (VOC) defining mutations in the SARS-CoV-2 spike gene.

| Lineage | a.a mutation in the spike of NY WTD SARS-CoV-2 | Proportion in VOC of NY WTD^1^ | VOC spike defining mutation |
| --- | --- | --- | --- |
| Alpha | D1118Y | 12.7% | D1118H |
| Alpha | C136S | 50% | D138H |
| Alpha | D568N | 5.5% | A570D |
| Alpha | I569T | 5.5% | A570D |
| Delta | R683W | 5.7% | P681R |

^1^ Proportion within VOC in sequences obtained in the present study

**Supplementary table 8.** Metadata table of samples used in the present study.

| **Short ID** | **GISAID accession number** | **Genbank accession number** | **Date** | **Country** | **State** | **County** | **Town** | **Cluster** | **Sex** | **Lineage classification** |
| --- | --- | --- | --- | --- | --- | --- | --- | --- | --- | --- |
| 119428 | EPI_ISL_13610627 | OP006287 | 2021-11-03 | USA | New York | STEU | Cohocton |  | M | Alpha |
| 119438 | EPI_ISL_13610615 | OP006288 | 2021-10-30 | USA | New York | ALLE | Friendship | C2 | M | Gamma |
| 119457 | EPI_ISL_13610654 | OP006289 | 2021-11-07 | USA | New York | LIVI | Dansville | C7 | M | Alpha |
| 119459 | EPI_ISL_13610602 | OP006290 | 2021-10-17 | USA | New York | LIVI | North Dansville | C7 | M | Alpha |
| 119471 | EPI_ISL_13610640 | OP006291 | 2021-11-06 | USA | New York | WYOM | Attica |  | M | Delta |
| 119478 | EPI_ISL_13610603 | OP006292 | 2021-10-18 | USA | New York | LIVI | Ossian | C7 | M | Alpha |
| 119480 | EPI_ISL_13610610 | OP006293 | 2021-10-27 | USA | New York | LIVI | Geneseo |  | M | Alpha |
| 119505 | EPI_ISL_13610625 | OP006294 | 2021-11-02 | USA | New York | ALLE | Burns | C7 | M | Alpha |
| 119521 | EPI_ISL_13610663 | OP006295 | 2021-11-09 | USA | New York | ALLE | Amity | C2 | M | Gamma |
| 119555 | EPI_ISL_13610611 | OP006296 | 2021-10-27 | USA | New York | ALLE | Belfast | C2 | M | Gamma |
| 119558 | EPI_ISL_13610621 | OP006297 | 2021-11-01 | USA | New York | SULL | Delaware | C1 | M | Delta |
| 119559 | EPI_ISL_13610655 | OP006298 | 2021-11-07 | USA | New York | ALLE | Wellsville | C2 | M | Gamma |
| 119567 | EPI_ISL_13610667 | OP006299 | 2021-11-11 | USA | New York | ORAN | Montgomery | C1 | M | Alpha |
| 119591 | EPI_ISL_13610613 | OP006300 | 2021-10-28 | USA | New York | SULL | Delaware | C1 | M | Delta |
| 119607 | EPI_ISL_13658779 | OP006301 | 2021-10-27 | USA | New York | ULST | Pine Bush | C1 | M | Alpha |
| 119611 | EPI_ISL_13610628 | OP006302 | 2021-11-03 | USA | New York | ORAN | Warwick | C1 | M | Delta |
| 119627 | EPI_ISL_13610678 | OP006303 | 2021-11-20 | USA | New York | ORAN | Montgomery | C1 | M | Alpha |
| 119639 | EPI_ISL_13610626 | OP006304 | 2021-11-02 | USA | New York | ALLE | Scio | C2 | M | Gamma |
| 119640 | EPI_ISL_13610605 | OP006305 | 2021-10-23 | USA | New York | ALLE | Alma | C2 | M | Gamma |
| 119642 | EPI_ISL_13610664 | OP006306 | 2021-11-09 | USA | New York | ALLE | Amity | C2 | M | Gamma |
| 119770 | EPI_ISL_13658805 | OP006307 | 2021-11-19 | USA | New York | DUTC | Verbank |  | M | Delta |
| 119777 | EPI_ISL_13658806 | OP006308 | 2021-11-20 | USA | New York | ULST | New Paltz |  | F | Alpha |
| 119785 | EPI_ISL_13610591 | OP006309 | 2021-10-02 | USA | New York | BROO | Conklin |  | F | Delta |
| 119867 | EPI_ISL_13610696 | OP006310 | 2021 | USA | New York | MADI | Sullivan |  | F | Alpha |
| 119941 | EPI_ISL_13658817 | OP006311 | 2021 | USA | New York | RENS |  |  | F | Alpha |
| 119945 | EPI_ISL_13610641 | OP006312 | 2021-11-06 | USA | New York | TOMP | Ithaca | C18 | M | Delta |
| 119983 | EPI_ISL_13610642 | OP006313 | 2021-11-06 | USA | New York | TOMP | Danby | C13 | M | Alpha |
| 119988 | EPI_ISL_13658807 | OP006314 | 2021-11-20 | USA | New York | CHEN | Sherburne |  | F | Alpha |
| 119997 | EPI_ISL_13610659 | OP006315 | 2021-11-08 | USA | New York | TIOG | Owego |  | M | Alpha |
| 140996 | EPI_ISL_13610693 | OP006316 | 2021-11-28 | USA | New York | SULL | Cochecton | C1 | F | Delta |
| 147551 | EPI_ISL_13610614 | OP006317 | 2021-10-29 | USA | New York | LEWI | Martinsburg | C8 | M | Delta |
| 149738 | EPI_ISL_13610691 | OP006318 | 2021-11-27 | USA | New York | TOMP | Ulysses | C18 | M | Delta |
| 149792 | EPI_ISL_13658808 | OP006319 | 2021-11-20 | USA | New York | OTSE | Roseboom |  | M | Delta |
| 149832 | EPI_ISL_13658813 | OP006320 | 2021-11-24 | USA | New York | ORAN | Hamptonburgh | C1 | M | Alpha |
| 149844 | EPI_ISL_13658816 | OP006321 | 2021-11-28 | USA | New York | SCHO | Jefferson |  | M | Delta |
| 149845 | EPI_ISL_13610688 | OP006322 | 2021-11-22 | USA | New York | ORAN | Vails Gate | C1 | M | Delta |
| 149849 | EPI_ISL_13610689 | OP006323 | 2021-11-22 | USA | New York | ORAN | Montgomery | C1 | M | Alpha |
| 149911 | EPI_ISL_13610599 | OP006324 | 2021-10-10 | USA | New York | RENS | East Greenbush |  | M | Alpha |
| 149932 | EPI_ISL_13610670 | OP006325 | 2021-11-13 | USA | New York | ONON | Manlius |  | M | Alpha |
| 149937 | EPI_ISL_13610588 | OP006326 | 2020-11-22 | USA | New York | MADI | Sullivan |  | M | Alpha |
| 149959 | EPI_ISL_13610697 | OP006327 | 2021 | USA | New York |  |  |  |  | Alpha |
| 149982 | EPI_ISL_13610671 | OP006328 | 2021-11-13 | USA | New York | CHAU | Portland |  | M | Delta |
| 149990 | EPI_ISL_13610668 | OP006329 | 2021-11-11 | USA | New York | CHAU | Charlotte |  | M | Delta |
| 152918 | EPI_ISL_13658777 | OP006330 | 2021-10-02 | USA | New York | ERIE | Orchard Park |  | M | Alpha |
| 154944 | EPI_ISL_13610593 | OP006331 | 2021-10-03 | USA | New York | ALBA | Voorheesville |  | M | Alpha |
| 155877 | EPI_ISL_13610592 | OP006332 | 2021-10-02 | USA | New York | LIVI | Portage | C7 | M | Alpha |
| 155912 | EPI_ISL_13658814 | OP006333 | 2021-11-26 | USA | New York | SULL | Callicoon | C1 | M | Delta |
| 155994 | EPI_ISL_13610616 | OP006334 | 2021-10-30 | USA | New York | CAYU | Aurelius |  | M | Delta |
| 158016 | EPI_ISL_13610619 | OP006335 | 2021-10-31 | USA | New York | ALLE | Wirt | C2 | M | Gamma |
| 158035 | EPI_ISL_13610677 | OP006336 | 2021-11-17 | USA | New York | CHAU | Carroll | C12 | M | Delta |
| 158072 | EPI_ISL_13610698 | OP006337 | 2021 | USA | New York | CATT | Mansfield |  | M | Delta |
| 158076 | EPI_ISL_13658818 | OP006338 | 2021 | USA | New York | CATT | Perrysburg |  | F | Alpha |
| 159048 | EPI_ISL_13610586 | OP006339 | 2020-11-13 | USA | New York | Sullivan | Liberty | C1 | M | B.1.409 |
| 159201 | EPI_ISL_13610584 | OP006340 | 2020-11-12 | USA | New York | Greene | Leeds |  | M | B.1.243 |
| 159212 | EPI_ISL_13610587 | OP006341 | 2020-11-21 | USA | New York | Ulster | Saugerties |  | M | B.1.243 |
| 161392 | EPI_ISL_13610582 | OP006342 | 2020-11-07 | USA | New York | Steuben | Addison |  | M | Delta |
| 161809 | EPI_ISL_13610699 | OP006343 | 2020 | USA | New York | Columbia | Ancram |  | F | B.1.517 |
| 161864 | EPI_ISL_13658781 | OP006344 | 2021-11-01 | USA | New York | ORAN | Chester | C1 | M | Delta |
| 161956 | EPI_ISL_13658787 | OP006345 | 2021-11-03 | USA | New York | ALLE | Amity | C2 | M | Gamma |
| 161965 | EPI_ISL_13610633 | OP006346 | 2021-11-05 | USA | New York | ALLE | Amity | C2 | M | Gamma |
| 162299 | EPI_ISL_13610580 | OP006347 | 2020-10-11 | USA | New York | Orleans | Carlton |  | F | B.1.1 |
| 162885 | EPI_ISL_13610581 | OP006348 | 2020-11-05 | USA | New York | Chemung | Horseheads |  | M | B.1.2 |
| 162900 | EPI_ISL_13610583 | OP006349 | 2020-11-07 | USA | New York | Monroe | Henrietta |  | M | B.1.1 |
| 162948 | EPI_ISL_13610600 | OP006350 | 2021-10-10 | USA | New York | Steuben | Campbell |  | M | B.1.2 |
| 162950 | EPI_ISL_13610585 | OP006351 | 2020-11-12 | USA | New York | Steuben | Lindley |  | M | B.1 |
| 163183 | EPI_ISL_13610675 | OP006352 | 2021-11-14 | USA | New York | ONEI | Trenton |  | M | Delta |
| 163188 | EPI_ISL_13658801 | OP006353 | 2021-11-10 | USA | New York | ONEI | Whitestown |  | M | Alpha |
| 163192 | EPI_ISL_13610672 | OP006354 | 2021-11-13 | USA | New York | ONEI | Westmoreland |  | M | Alpha |
| 164040 | EPI_ISL_13610597 | OP006355 | 2021-10-09 | USA | New York | CHAU | North Harmony | C12 | M | Delta |
| 164122 | EPI_ISL_13610601 | OP006356 | 2021-10-13 | USA | New York | LIVI | Sparta | C7 | F | Alpha |
| 164135 | EPI_ISL_13610598 | OP006357 | 2021-10-09 | USA | New York | ORLE | Ridgeway |  | F | Delta |
| 164175 | EPI_ISL_13610604 | OP006358 | 2021-10-19 | USA | New York | GENE | Pembroke |  | M | Delta |
| 164255 | EPI_ISL_13658780 | OP006359 | 2021-10-30 | USA | New York | ERIE | Aurora |  | F | Alpha |
| 164331 | EPI_ISL_13658778 | OP006360 | 2021-10-23 | USA | New York | CHAU | Busti | C12 | M | Alpha |
| 164343 | EPI_ISL_13610606 | OP006361 | 2021-10-23 | USA | New York | CHAU | North Harmony | C12 | F | Delta |
| 164345 | EPI_ISL_13610607 | OP006362 | 2021-10-24 | USA | New York | ALLE | Cuba | C2 | M | Gamma |
| 164384 | EPI_ISL_13610617 | OP006363 | 2021-10-30 | USA | New York | ALLE | Wellsville | C2 | F | Alpha |
| 164389 | EPI_ISL_13610612 | OP006364 | 2021-10-27 | USA | New York | ALLE | Bolivar | C2 | F | Gamma |
| 164395 | EPI_ISL_13610608 | OP006365 | 2021-10-24 | USA | New York | WAYN | Butler |  | M | Delta |
| 164412 | EPI_ISL_13610622 | OP006366 | 2021-11-01 | USA | New York | CHAU | Poland | C12 | M | Alpha |
| 164447 | EPI_ISL_13658782 | OP006367 | 2021-11-02 | USA | New York | ERIE | Newstead |  | F | Delta |
| 164453 | EPI_ISL_13610643 | OP006368 | 2021-11-06 | USA | New York | ALLE | Wirt | C2 | M | Gamma |
| 164458 | EPI_ISL_13610618 | OP006369 | 2021-10-30 | USA | New York | TIOG | Spencer | C13 | M | Alpha |
| 164463 | EPI_ISL_13610620 | OP006370 | 2021-10-31 | USA | New York | ALLE | Wirt | C2 | M | Gamma |
| 164465 | EPI_ISL_13610644 | OP006371 | 2021-11-06 | USA | New York | ALLE | Bolivar | C2 | M | Gamma |
| 164468 | EPI_ISL_13658792 | OP006372 | 2021-11-06 | USA | New York | CHAU | Busti | C12 | M | Alpha |
| 164472 | EPI_ISL_13658788 | OP006373 | 2021-11-03 | USA | New York | CHAU | North Harmony | C12 | M | Alpha |
| 164477 | EPI_ISL_13610656 | OP006374 | 2021-11-07 | USA | New York | CHAU | Chautauqua |  | F | Delta |
| 164485 | EPI_ISL_13610629 | OP006375 | 2021-11-03 | USA | New York | CHAU | Clymer |  | M | Delta |
| 164487 | EPI_ISL_13610630 | OP006376 | 2021-11-03 | USA | New York | CATT | Randolph | C12 | M | Alpha |
| 164489 | EPI_ISL_13658793 | OP006377 | 2021-11-06 | USA | New York | CHAU | Kiantone | C12 | M | Alpha |
| 164493 | EPI_ISL_13610594 | OP006378 | 2021-10-04 | USA | New York | ALLE | Cuba | C2 | M | Gamma |
| 164507 | EPI_ISL_13658794 | OP006379 | 2021-11-06 | USA | New York | CHAU | Busti | C12 | F | Delta |
| 164509 | EPI_ISL_13658799 | OP006380 | 2021-11-07 | USA | New York | CATT | East Otto |  | M | Delta |
| 164510 | EPI_ISL_13658795 | OP006284 | 2021-11-06 | USA | New York | CHAU | Busti | C12 | M | Delta |
| 164522 | EPI_ISL_13610634 | OP006381 | 2021-11-05 | USA | New York | ERIE | Aurora |  | M | Alpha |
| 164523 | EPI_ISL_13610635 | OP006382 | 2021-11-05 | USA | New York | CHAU | Harmony | C12 | F | Delta |
| 164525 | EPI_ISL_13610595 | OP006383 | 2021-10-04 | USA | New York | CHAU | Portland |  | M | Delta |
| 164529 | EPI_ISL_13658783 | OP006384 | 2021-11-02 | USA | New York | ALLE | Amity | C2 | F | Gamma |
| 164530 | EPI_ISL_13610636 | OP006385 | 2021-11-05 | USA | New York | CHAU | Chautauqua |  | M | Delta |
| 164531 | EPI_ISL_13610589 | OP006386 | 2021-10-01 | USA | New York | CHAU | Chautauqua |  | M | Delta |
| 164533 | EPI_ISL_13658800 | OP006387 | 2021-11-07 | USA | New York | MONR | Rush |  | M | Alpha |
| 164544 | EPI_ISL_13610645 | OP006388 | 2021-11-06 | USA | New York | ALLE | Angelica |  | M | Gamma |
| 164548 | EPI_ISL_13610657 | OP006389 | 2021-11-07 | USA | New York | ALLE | Willing | C2 | M | Gamma |
| 164550 | EPI_ISL_13610590 | OP006390 | 2021-10-01 | USA | New York | CHAU | Chautauqua |  | M | Delta |
| 164553 | EPI_ISL_13610660 | OP006391 | 2021-11-08 | USA | New York | ALLE | Wirt | C2 | M | Gamma |
| 164557 | EPI_ISL_13658796 | OP006392 | 2021-11-06 | USA | New York | CHAU | Busti | C12 | M | Alpha |
| 164565 | EPI_ISL_13610623 | OP006393 | 2021-11-01 | USA | New York | STEU | Greenwood |  | M | Gamma |
| 164571 | EPI_ISL_13658789 | OP006394 | 2021-11-04 | USA | New York | TOMP | Enfield | C18 | M | Delta |
| 164572 | EPI_ISL_13610596 | OP006395 | 2021-10-08 | USA | New York | WYOM | Warsaw |  | F | Delta |
| 164573 | EPI_ISL_13658784 | OP006396 | 2021-11-02 | USA | New York | CHAU | French Creek |  | M | Delta |
| 164580 | EPI_ISL_13610646 | OP006397 | 2021-11-06 | USA | New York | CHAU | Chautauqua |  | F | Delta |
| 164604 | EPI_ISL_13610647 | OP006398 | 2021-11-06 | USA | New York | WAYN | Galen |  | F | Delta |
| 164628 | EPI_ISL_13610700 | OP006399 | 2021 | USA | New York | WYOM | Bennington |  | F | Delta |
| 164637 | EPI_ISL_13610648 | OP006400 | 2021-11-06 | USA | New York | TOMP | Danby | C13 | M | Delta |
| 164656 | EPI_ISL_13610631 | OP006401 | 2021-11-03 | USA | New York | CHAU | Westfield |  | M | Delta |
| 164661 | EPI_ISL_13610649 | OP006402 | 2021-11-06 | USA | New York | ALLE | Amity | C2 | F | Gamma |
| 164669 | EPI_ISL_13610632 | OP006403 | 2021-11-03 | USA | New York | ALLE | Alma | C2 | M | Gamma |
| 164672 | EPI_ISL_13610637 | OP006404 | 2021-11-05 | USA | New York | ALLE | Cuba | C2 | M | Gamma |
| 164673 | EPI_ISL_13658797 | OP006405 | 2021-11-06 | USA | New York | ALLE | Cuba | C2 | M | Gamma |
| 164676 | EPI_ISL_13610673 | OP006406 | 2021-11-13 | USA | New York | ALLE | Scio | C2 | M | Gamma |
| 164702 | EPI_ISL_13610661 | OP006407 | 2021-11-08 | USA | New York | CHEM | Baldwin | C13 | M | Delta |
| 164706 | EPI_ISL_13610638 | OP006408 | 2021-11-05 | USA | New York | GENE |  |  | M | Delta |
| 164707 | EPI_ISL_13610650 | OP006409 | 2021-11-06 | USA | New York | WYOM | Pike | C7 | M | Alpha |
| 164709 | EPI_ISL_13610651 | OP006410 | 2021-11-06 | USA | New York | CATT | New Albion |  | M | Delta |
| 164713 | EPI_ISL_13610676 | OP006411 | 2021-11-15 | USA | New York | TIOG | Spencer | C13 | M | Alpha |
| 164717 | EPI_ISL_13658804 | OP006285 | 2021-11-15 | USA | New York | LIVI | Avon |  | M | Alpha |
| 164723 | EPI_ISL_13658790 | OP006412 | 2021-11-05 | USA | New York | ALLE | Hume | C7 | M | Alpha |
| 164729 | EPI_ISL_13658798 | OP006413 | 2021-11-06 | USA | New York | ALLE | Amity | C2 | M | Gamma |
| 164730 | EPI_ISL_13610695 | OP006414 | 2021-12-27 | USA | New York | ALLE | Granger | C7 | M | Alpha |
| 164733 | EPI_ISL_13658803 | OP006415 | 2021-11-12 | USA | New York | TOMP | Ulysses | C18 | M | Delta |
| 164737 | EPI_ISL_13610652 | OP006416 | 2021-11-06 | USA | New York | CATT | Farmersville |  | M | Alpha |
| 164739 | EPI_ISL_13658802 | OP006286 | 2021-11-11 | USA | New York | ALLE | Centerville |  | M | Alpha |
| 164750 | EPI_ISL_13610658 | OP006418 | 2021-11-07 | USA | New York | WYOM | Perry |  | M | B.1.2 |
| 164755 | EPI_ISL_13610639 | OP006419 | 2021-11-05 | USA | New York | WYOM | Pike | C7 | M | Alpha |
| 164760 | EPI_ISL_13610653 | OP006420 | 2021-11-06 | USA | New York | CATT | Yorkshire |  | M | Delta |
| 164770 | EPI_ISL_13610674 | OP006421 | 2021-11-13 | USA | New York | ALLE | Caneadea |  | F | B.1.2 |
| 164777 | EPI_ISL_13610665 | OP006422 | 2021-11-09 | USA | New York | TOMP | Ulysses | C18 | F | Delta |
| 164781 | EPI_ISL_13610662 | OP006423 | 2021-11-08 | USA | New York | CHEM | Baldwin | C13 | F | B.1.1 |
| 164784 | EPI_ISL_13610669 | OP006424 | 2021-11-12 | USA | New York | CHAU | Bemus Point |  | M | Delta |
| 165189 | EPI_ISL_13658791 | OP006417 | 2021-11-05 | USA | New York | SUFF | Southold |  | M | Delta |
| 165273 | EPI_ISL_13610666 | OP006425 | 2021-11-10 | USA | New York | WASH | Fort Ann |  | M | Delta |
| 165621 | EPI_ISL_13658785 | OP006426 | 2021-11-02 | USA | New York | TIOG | Spencer | C13 | M | Alpha |
| 165628 | EPI_ISL_13610687 | OP006427 | 2021-11-21 | USA | New York | TIOG | Berkshire |  | M | Alpha |
| 165672 | EPI_ISL_13658811 | OP006428 | 2021-11-21 | USA | New York | DUTC | Amenia |  | M | Delta |
| 165681 | EPI_ISL_13658809 | OP006429 | 2021-11-20 | USA | New York | SULL | Bloomingburg | C1 | F | Alpha |
| 165697 | EPI_ISL_13610679 | OP006430 | 2021-11-20 | USA | New York | SULL | Delaware | C1 | F | Delta |
| 165707 | EPI_ISL_13610680 | OP006431 | 2021-11-20 | USA | New York | DUTC | Washington |  | M | Delta |
| 165711 | EPI_ISL_13610681 | OP006432 | 2021-11-20 | USA | New York | SULL | Tusten | C1 | M | Delta |
| 165715 | EPI_ISL_13610682 | OP006433 | 2021-11-20 | USA | New York | SULL | Tusten | C1 | M | Delta |
| 165724 | EPI_ISL_13610692 | OP006434 | 2021-11-27 | USA | New York | ORAN | Chester | C1 | F | Delta |
| 165726 | EPI_ISL_13658810 | OP006435 | 2021-11-20 | USA | New York | SULL | Cochecton | C1 | F | Delta |
| 165733 | EPI_ISL_13610683 | OP006436 | 2021-11-20 | USA | New York | ONON | Cicero |  | F | Alpha |
| 165741 | EPI_ISL_13610690 | OP006437 | 2021-11-26 | USA | New York | SULL | Thompson | C1 | F | Delta |
| 165751 | EPI_ISL_13610701 | OP006438 | 2021 | USA | New York | SULL | Delaware | C1 | M | Delta |
| 165776 | EPI_ISL_13610684 | OP006439 | 2021-11-20 | USA | New York | WAYN | Butler |  | M | Delta |
| 165804 | EPI_ISL_13658812 | OP006440 | 2021-11-22 | USA | New York | ALBA | Medusa |  | M | Delta |
| 165834 | EPI_ISL_13610685 | OP006441 | 2021-11-20 | USA | New York | TIOG | Newark Valley |  | M | Delta |
| 165887 | EPI_ISL_13610624 | OP006442 | 2021-11-01 | USA | New York | ONON | Camillus |  | M | Delta |
| 165892 | EPI_ISL_13610686 | OP006443 | 2021-11-20 | USA | New York | TIOG | Owego |  | M | Alpha |
| 165908 | EPI_ISL_13610702 | OP006444 | 2021 | USA | New York |  |  |  |  | Alpha |
| 165920 | EPI_ISL_13610694 | OP006445 | 2021-12-04 | USA | New York | TIOG | Owego |  | F | Alpha |
| 165925 | EPI_ISL_13658815 | OP006446 | 2021-11-27 | USA | New York | CAYU | Throop |  | F | Delta |
| 165954 | EPI_ISL_13610609 | OP006447 | 2021-10-24 | USA | New York | CHEN | Greene |  | M | Delta |
